# Endosomal free fatty acid receptor 2 signaling is essential for propionate-induced anorectic gut hormone release

**DOI:** 10.1101/2020.03.24.004762

**Authors:** Natarin Caengprasath, Noemi Gonzalez-Abuin, Maria Shchepinova, Yue Ma, Asuka Inoue, Edward W. Tate, Gary Frost, Aylin C. Hanyaloglu

**Affiliations:** Institute of Reproductive and Developmental Biology, Department of Metabolism, Digestion and Reproduction, Imperial College London, London, UK; Department of Metabolism, Digestion and Reproduction, Imperial College London, London, UK; Department of Chemistry, Imperial College London, London, UK; Graduate School of Pharmaceutical Sciences, Tohoku University, Sendai, Japan

## Abstract

The ability of propionate, a short chain fatty acid produced from the fermentation of non-digestible carbohydrates in the colon, to stimulate the release of anorectic gut hormones, such as glucagon like peptide-1 (GLP-1), is an attractive approach to enhance appetite regulation, weight management and glycaemic control. Propionate induces GLP-1 release via its G protein-coupled receptor (GPCR), free fatty acid receptor 2 (FFA2); a GPCR that activates Gαi and Gαq/11 pathways. However, how pleiotropic GPCR signaling mechanisms in the gut regulates appetite is poorly understood. Here, we identify propionate-mediated G protein signaling is spatially directed within the cell via the targeting of FFA2 to very early endosomes. Furthermore, propionate activates an endosomal Gαi/p38 signaling pathway, which is essential for propionate-induced GLP-1 release in enteroendocrine cells and colonic crypts. Our study reveals that intestinal metabolites can engage membrane trafficking pathways and endosomal signaling platforms to orchestrate complex GPCR pathways within the gut.

## Introduction

The consumption of dietary fiber, or non-digestible carbohydrates (NDCs), has been shown to protect against diet-induced obesity (Chambers et al., 2015). The protective effects of NDCs are largely attributed to short chain fatty acids (SCFAs) that are produced in the colon by microbiota from the fermentation of NDCs (Chambers et al., 2015; den Besten et al., 2013; James et al., 2003). Acetate, propionate and butyrate are the predominant SCFAs produced and in addition to regulation of gastro-intestinal functions, are involved in energy and glucose homeostasis and immune responses (den Besten et al., 2013). Traditionally, roles of SCFAs in these metabolic processes were thought to be limited to their ability to act as an energy source or as a regulator of cholesterol synthesis, however, with the discovery and characterization of G protein-coupled receptors (GPCRs) activated by SCFAs, free fatty acid receptor 2 (FFA2, previously known as GPR43) and free fatty acid receptor 3 (FFA3, previously known as GPR41), it is now widely appreciated that many SCFA activities can be attributed to these receptors (Fuller et al., 2015; Li et al., 2018; Pingitore et al., 2019; Tolhurst et al., 2012; Bolognini et al., 2016).

Among the three SCFAs, propionate has been of particular translational interest due to its ability to acutely suppress appetite via activation of FFA2 in enteroendocrine L cells, and release of the anorectic gut hormones peptide YY (PYY) and incretin glucagon like peptide-1 (GLP-1) (Tolhurst et al., 2012; Psichas et al., 2015), contributing to its role in rapid weight loss and improved insulin sensitivity following roux-en-Y gastric bypass (Liou et al., 2013). Direct health benefits of propionate in humans have been recently demonstrated whereby increasing the colonic levels of propionate in overweight humans not only exhibited reduced weight gain, but also reduced abdominal adiposity and improved insulin sensitivity (Chambers et al., 2015). Thus propionate, and its receptor-mediated actions, represent an attractive system to develop therapeutic strategies in obesity management.

Although the role of SCFAs and their receptors in mediating the release of anorectic gut hormones has been demonstrated in rodent models and humans (Tolhurst et al., 2012; Bolognini et al., 2016; Psichas et al., 2015; Chambers et al., 2015), our understanding of the molecular mechanisms by propionate, regulates the release of anorectic gut hormone from enteroendocrine L cells remains limited. FFA2 is coupled to both the Gαi/o and Gαq/11 families of heterotrimeric G proteins (Brown et al., 2003; Le Poul et al., 2003), although Gαq/11 is implicated in mediating gut hormone release via increases in calcium (Bolognini et al., 2016; Tolhurst et al., 2012). Models of GPCR signaling, however, have rapidly evolved over recent years from single receptors activating distinct G protein pathways at the plasma membrane, to high signal diversity that can be differentially activated by distinct ligands and exquisitely regulated at a spatial and temporal level. The spatio-temporal regulation of GPCRs can occur via a variety of processes, with membrane trafficking of GPCRs playing a central role. Membrane trafficking of GPCRs was classically viewed as a mechanism to control active cell surface receptor number by driving receptor internalization and post-endocytic sorting to divergent cellular fates. However, it is now understood that receptor internalization to endosomes provides additional intracellular signaling platforms including activation of heterotrimeric G protein signaling (Eichel and von Zastrow, 2018; Hanyaloglu, 2018). Endosomal signaling of GPCRs exhibits distinct functions from signaling activated at the plasma membrane, demonstrating the integrated nature of trafficking and signaling, and providing a mechanism for cells to achieve highly-specific and diverse downstream responses to its dynamic extracellular environment (Thomsen et al., 2018; Caengprasath and Hanyaloglu, 2019). Furthermore, we have previously shown that GPCRs are organized to distinct endosomal compartments to activate signaling (Sposini et al., 2017; Jean-Alphonse et al., 2014). These discoveries over the past decade have rewritten the GPCR ‘signaling atlas’, offering new interpretations of faulty GPCR activity in disease and providing novel therapeutic strategies to target GPCR signaling (Thomsen et al., 2018). However, the role of membrane trafficking for FFA2, and the distinct actions of propionate that activates pleiotropic G protein signal pathways within the gut, remain unknown.

In this study, we demonstrate a key role for endosomal FFA2 signaling to drive propionate-induced GLP-1 release from enteroendocrine cells. Furthermore, we provide evidence that G protein signaling activated by FFA2 is differentially regulated by membrane trafficking, and that an unexpected endosomal Gαi/p38 signaling pathway is required for propionate-induced GLP-1 release.

## Results

### Propionate stimulates GLP-1 secretion yet activates Gαi/o but not Gαq/11 signaling from colonic crypts and enteroendocrine cells

Although propionate is known to mediate anorectic gut hormone release via FFA2, the ability of this SCFA to activate both upstream Gαq/11 and Gαi/o signal pathways in enteroendocrine cells has yet to be fully demonstrated. FFA2 couples to both Gαi/o to inhibit adenylate cyclase and reduce intracellular levels of cAMP, and Gαq/11 that activates phospholipase C resulting in increases in inositol 1,4,5-triphosphate (IP_3_) and diacylglycerol, leading to mobilization of calcium from intracellular stores.

In both mouse enteroendocrine (STC-1) cells and colonic crypts, propionate was able to inhibit forskolin-induced cAMP, which was significantly reversed by pre-treatment with Gαi/o inhibitor pertussis toxin (Ptx) (Figure 1A and 1B). Surprisingly, propionate did not induce either an increase in intracellular calcium (Figure 1C and 1D) or IP_1_, a downstream metabolite of IP_3_, in either STC-1 cells or colonic crypts (Figure 1E and 1F) despite its ability to induce GLP-1 release (Figure 1G and 1H). In contrast, a previously described selective FFA2 synthetic allosteric ligand (Lee et al., 2008) 4-CTMB, and a previously characterized selective FFA2 synthetic orthosteric ligand, compound 1 (Hudson et al., 2013) (Cmp1) activated both Gαi/o and Gαq/11 signaling in STC-1 cells and colonic crypts (Figure S1A, 1C-1F). Thus, further demonstrating functional FFA2 in both cultures. The synthetic ligand-induced calcium responses were Gαq/11-mediated as they were significantly impaired by the pre-treatment of a selective Gαq/11 inhibitor, YM-254890 (Takasaki et al., 2004) in STC-1 cells (Figure S1B).

**Figure 1.**
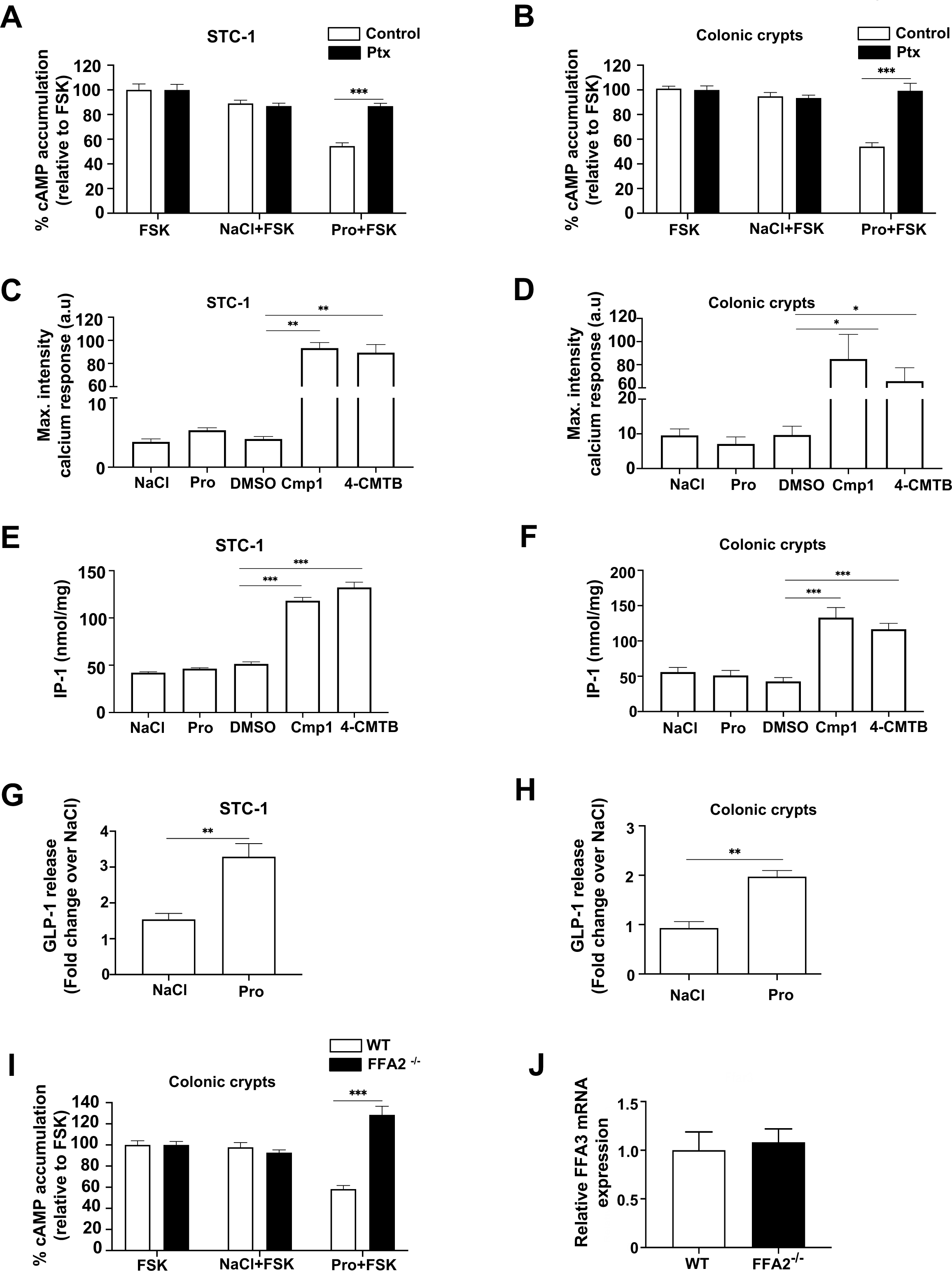
Propionate stimulates GLP-1 secretion and activates Gαi/o but not Gαq/11 via FFA2. (A-B) Intracellular cAMP levels measured in STC-1 cells (A) or colonic crypts (B) pre-treated with Pertussis toxin (Ptx; 200ng/mL, 20h) prior to pre-treatment with IBMX (0.5 mM, 5 min) and then stimulated with forskolin (FSK, 3 µM) or a combination of FSK with either NaCl or sodium propionate (Pro) (1 mM, 5 min). Data are expressed as % change of FSK treated cells. n = 3 independent experiments. Two-sided Mann-Whitney U test, *** p < 0.001. (C-D) Intracellular calcium mobilization measured in STC-1 cells (C) or colonic crypts (D). Cultures were incubated with calcium indicator Fluo4-AM for 1 h and imaged live via confocal microscopy for 1 min before the addition of either NaCl (1 mM), sodium propionate (Pro), DMSO, orthosteric FFA2 agonist Cmp1 (10 µM) or allosteric FFA2 agonist 4-CMTB (10 µM). Average maximal intensities of n = 20 cells in duplicate per 6 independent experiments. Two-sided Mann-Whitney U test, * p < 0.05. (E-F) Intracellular accumulation of IP_1_ in STC-1 cells (E) or colonic crypts (F). Cultures were treated with either NaCl, sodium propionate (Pro) (1 mM), DMSO, Cmp1 (10 μM) or 4-CMTB (10 μM) for 2 h. STC-1 cells or crypts, n = 3 independent experiments. Two-sided Mann-Whitney U test, *** p < 0.001. Data represent mean ± SEM. (G) STC-1 cells and (H) colonic crypts were treated with either NaCl, sodium propionate (Pro) (1 mM, 2 h STC-1, 1 h crypts) and total GLP-1 levels secreted was measured via RIA. Data are expressed as fold change of total GLP-1 and normalized to basal (NaCl) secretion within the same experiment. For STC-1 cells, n = 4 independent experiments. For crypts, n = 3 independent experiments. Two-sided Mann-Whitney U test, * p < 0.05, ** p < 0.01. (I) Intracellular cAMP levels measured in colonic crypts from wildtype (WT) or FFA2 knockout mice (FFA2 ^-/-^) pre-treated with IBMX (0.5 mM, 5 min) and then stimulated as in (C-D). Data are expressed as % change of FSK/NaCl treated cells. n = 3 independent experiments. Two-sided Mann-Whitney U test, *** p < 0.001. (J) Expression levels of FFA3 in WT and FFA2 ^-/-^ colonic crypts. mRNA isolated from colonic crypts of WT and FFA2 ^-/-^ mice were used in qPCR studies with specific mouse FFA3 primers. Data are presented as ΔΔCt. Two-sided Mann-Whitney U test. Data represent mean ± SEM.

Despite the lack of detectable Gαq/11 signaling, propionate-induced-Gαi/o signaling was dependent on FFA2 as the reduction of forskolin-induced cAMP in colonic crypts derived from FFA2 knockout mice (FFA2 ^-/-^) was completely abolished (Figure 1I), consistent with our prior reports from the same mouse model that propionate-induced gut hormone release from the colon requires FFA2 (Psichas et al., 2015). This loss of propionate-mediated Gαi/o signaling in the FFA2 ^-/-^ crypts was not due to alterations in Ffar3 expression (Figure 1J).

These data confirm that despite functional FFA2 responses, propionate activates Gαi/o signaling without detectable Gαq/11 responses in these cultures. To determine if the inability of propionate to activate Gαq/11 signaling via FFA2 was cellular context-specific, we stimulated HEK 293 cells expressing FLAG-FFA2. Upon stimulation with SCFAs, propionate significantly induced increases in intracellular calcium and IP_1_, (Figure S1C and S1D) confirming activation of Gαq/11 signaling. Taken together, this demonstrates that unlike synthetic FFA2 ligands, propionate is not able to signal via Gαq/11 in enteroendocrine cells, suggesting additional mechanisms beyond G-protein activation are employed to induce propionate-mediated anorectic gut hormone secretion.

### FFA2/G protein signaling is spatially regulated

We next determined if propionate/FFA2 activation is spatially regulated via membrane trafficking as a potential mechanism underlying its actions in the gut. Many GPCRs undergo ligand-induced internalization via a well described β-arrestin- and clathrin-dependent mechanism, whereby the large GTPase dynamin regulates the latter steps of endocytosis that drive clathrin-coated vesicle scission. To inhibit FFA2 internalization the ability of a potent inhibitor of dynamin GTPase activity, Dyngo-4a, known to block the internalization of many GPCRs (McCluskey et al., 2013; Eichel et al., 2016; Tsvetanova and von Zastrow, 2014; Sposini et al., 2017), was first assessed in HEK 293 cells expressing FLAG-tagged FFA2 and imaged via confocal microscopy. Unexpectedly, FFA2 exhibited both constitutive and propionate-dependent internalization from the plasma membrane (Figure 2A), which was confirmed via flow cytometry (Figure S2A). In cells pre-treated with Dyngo-4a, a strong inhibition of both constitutive and propionate-induced FFA2 internalization was observed (Figure 2A), demonstrating that FFA2 constitutive and ligand-induced internalization occur in a dynamin-dependent manner. Under conditions where dynamin-dependent FFA2 internalization was inhibited in HEK 293 cells, the ability of propionate to inhibit forskolin-induced cAMP was impaired (Figure 2B). In contrast, FFA2-mediated Gαq/11 signaling, as measured by intracellular calcium responses (Figure 2C) or IP-1 accumulation (Figure S2B), was not significantly affected, suggesting a differential requirement of FFA2 internalization for FFA2 mediated signaling. These results were also confirmed in HEK 293 cells lacking β-arrestins 1 and 2 (Grundmann et al., 2018) (Figure S2C). Interestingly, only ligand-induced, but not constitutive, FFA2 internalization was inhibited by lack of β-arrestins (Figure S2D). However, as in cells pre-treated with Dyngo-4a, propionate-dependent inhibition of forskolin-induced cAMP was significantly impaired in β-arrestin knockout cells compared to wildtype HEK 293 cells (Figure 2E). In contrast, propionate-induced calcium mobilization and IP_1_ accumulation was unperturbed (Figure 2F and Figure S2E).

**Figure 2.**
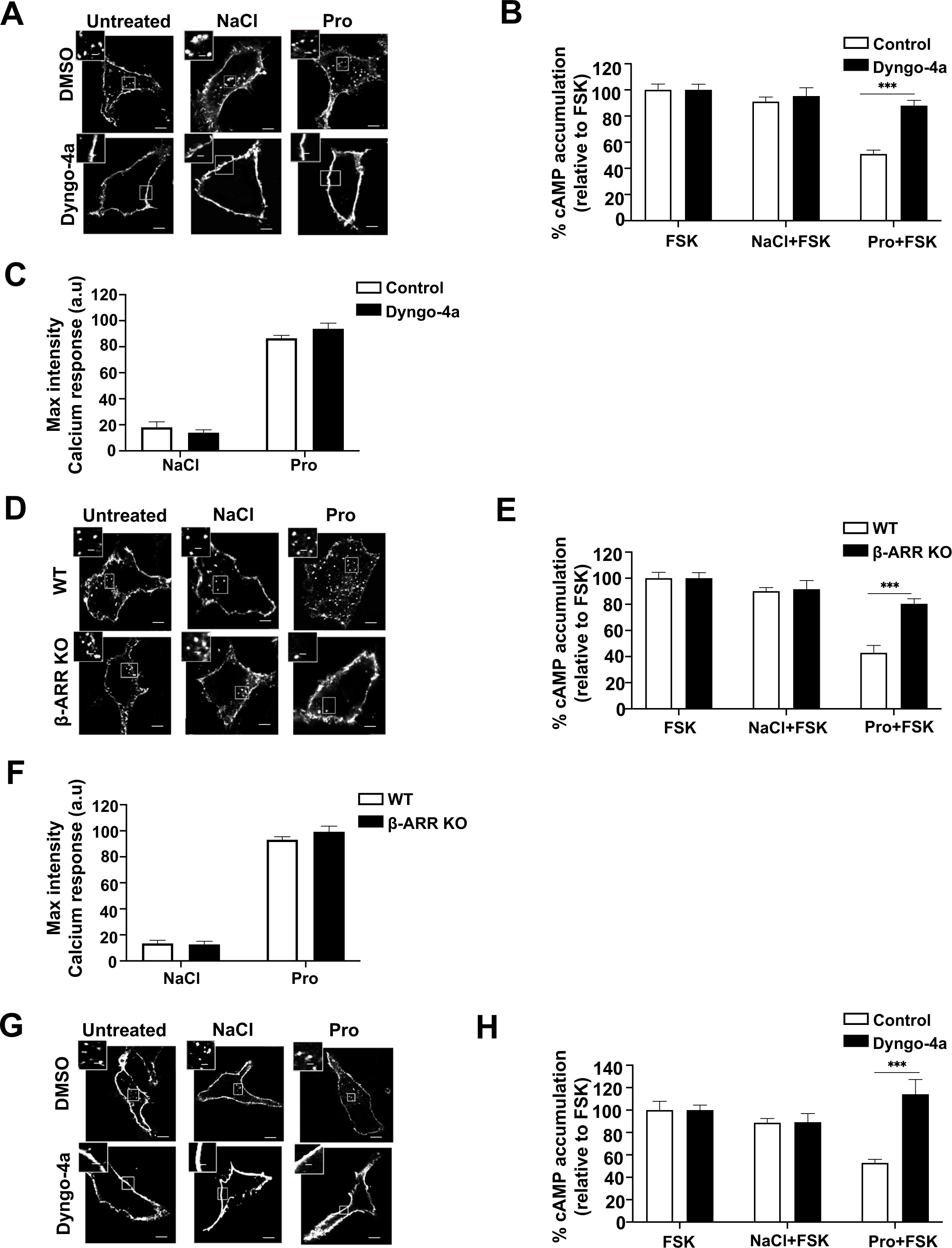
Propionate-dependent Gαi/o signaling requires receptor internalization. (A) Representative confocal microscopy images of HEK 293 cells expressing FLAG-FFA2 were pre-treated with either DMSO (vehicle) or Dyngo-4a (50 µM, 45 min), fed with M1 anti-FLAG antibody prior to stimulation with either NaCl or sodium propionate (Pro) (1 mM, 20 min). Fixed cells were imaged via confocal microscopy. (B-C) Intracellular cAMP levels (B) or calcium mobilization (C) measured in HEK 293 cells expressing FLAG-FFA2 pre-treated with either DMSO (vehicle) or Dyngo-4a (50 µM, 45 min). For (B), cells were pre-treated with IBMX (0.5 mM, 5 min) and then stimulated with forskolin (FSK, 3 µM) or a combination of FSK with either NaCl or sodium propionate (Pro) (1 mM, 5 min). n = 3 independent experiments. Two-sided Mann-Whitney U test, *** p < 0.001 For (C), cells were incubated with calcium indicator Fluo4-AM for 1 h and imaged live via confocal microscopy for 1 min before the addition of either NaCl or sodium propionate (Pro) (1 mM). Average maximal intensities of n=20 cells in duplicate per 4 independent experiments. (D) Representative confocal microscopy images of WT or β-ARR KO HEK 293 cells expressing FLAG-FFA2. Cells were treated with FLAG antibody and ligands and imaged as in (A). (E-F) Intracellular cAMP levels (E) or calcium mobilization (F) measured in WT or β-ARR KO HEK 293 cells transiently expressing FLAG-FFA2. Samples were treated and assayed as in (B) and (C). n = 3 independent experiments for either WT or β-ARR KO HEK 293 cells transiently expressing FLAG-FFA2. Two-sided Mann-Whitney U test, *** p < 0.001 (G) Representative confocal images of STC-1 cells transiently expressing FLAG-FFA2 pre-treated with either DMSO (vehicle) or Dyngo-4a (50 µM, 45 min) then stimulated as in (A). (H) Intracellular cAMP levels of STC-1 pre-treated with either DMSO (vehicle) or Dyngo-4a (50 µM, 45 min). Scale bar, 5 µm; scale bar in inset, 1 µm. n = 3 independent experiments. Two-sided Mann-Whitney U test, *** p < 0.001. For confocal images, representative images are shown of ∼10 cells/experiment. Data represent mean ± SEM.

The requirement of receptor internalization for Gα_i/o_ signaling was also determined for the endogenous propionate-responsive receptors expressed in STC-1 cells. As specific antibodies are not available for these receptors, STC-1 cells were transfected with FLAG-tagged FFA2 to confirm required conditions to inhibit FFA2 internalization in these cells. Similar, to HEK 293 cells, FFA2 internalization exhibited both constitutive and ligand-induced endocytic profiles, and both were inhibited by Dyngo-4a (Figure 2G). Consistent with our observations in HEK 293 cells, Dyngo-4a pre-treatment inhibited propionate-mediated activation of Gαi/o signaling (Figure 2H). Overall, these data demonstrate a requirement for ligand-induced FFA2 internalization in propionate-mediated Gαi/o signaling in heterologous and enteroendocrine cells.

### FFA2 internalizes to very early endosomes for sorting and signaling

We have previously shown that GPCRs exhibit divergent sorting to distinct endosomal compartments between early endosomes (EEs) and very early endosomes (VEEs), and this post-endocytic organization is critical for both GPCR sorting fate and endosomal signaling (Jean-Alphonse et al., 2014; Sposini et al., 2017). As internalization of FFA2 is essential for its Gαi/o signaling, we next determined the postendocytic compartment that FFA2 internalizes to. The organization of FFA2 across VEEs and EEs was compared with the β2-adrenergic receptor (β2AR), a GPCR known to be rapidly sorted to the EE compartment (Jean-Alphonse et al., 2014). VEEs are one third the diameter of EEs, and lack classic EE markers such as early endosomal autoantigen 1 (EEA1); however, a subpopulation of VEEs contain the adaption protein APPL1 (adaptor protein phosphotyrosine interaction, pleckstrin homology domain and, leucine zipper containing 1), which plays essential roles in post-VEE GPCR sorting and in regulating G protein signaling from this compartment (Jean-Alphonse et al., 2014; Sposini et al., 2017).

Internalization of FLAG-FFA2 was imaged in both live HEK 293 and STC-1 cells. FFA2 internalized to ∼400 nm diameter endosomes (Figure 3A), in contrast to the significantly larger size of endosomes containing internalized β2AR (Jean-Alphonse et al., 2014) (Figure 3A). Furthermore, the majority (>60%) of constitutive and ligand-induced internalized FFA2 did not traffic to an EEA1 positive EE compartment, compared to ∼70% for β2AR that did localize to EEA1 positive endosomes (Figure 3B), suggesting that FFA2 may traffic primarily to VEEs than EEs. This was further supported by the finding that a subpopulation of internalized FFA2 co-localizes with APPL1, (32.8 ± 0.35% following propionate treatment), similar to that observed with the luteinizing hormone receptor (LHR; 35.2 ± 1.92%), a GPCR known to traffic to VEEs (Sposini et al., 2017) (Figure 3C).

**Figure 3.**
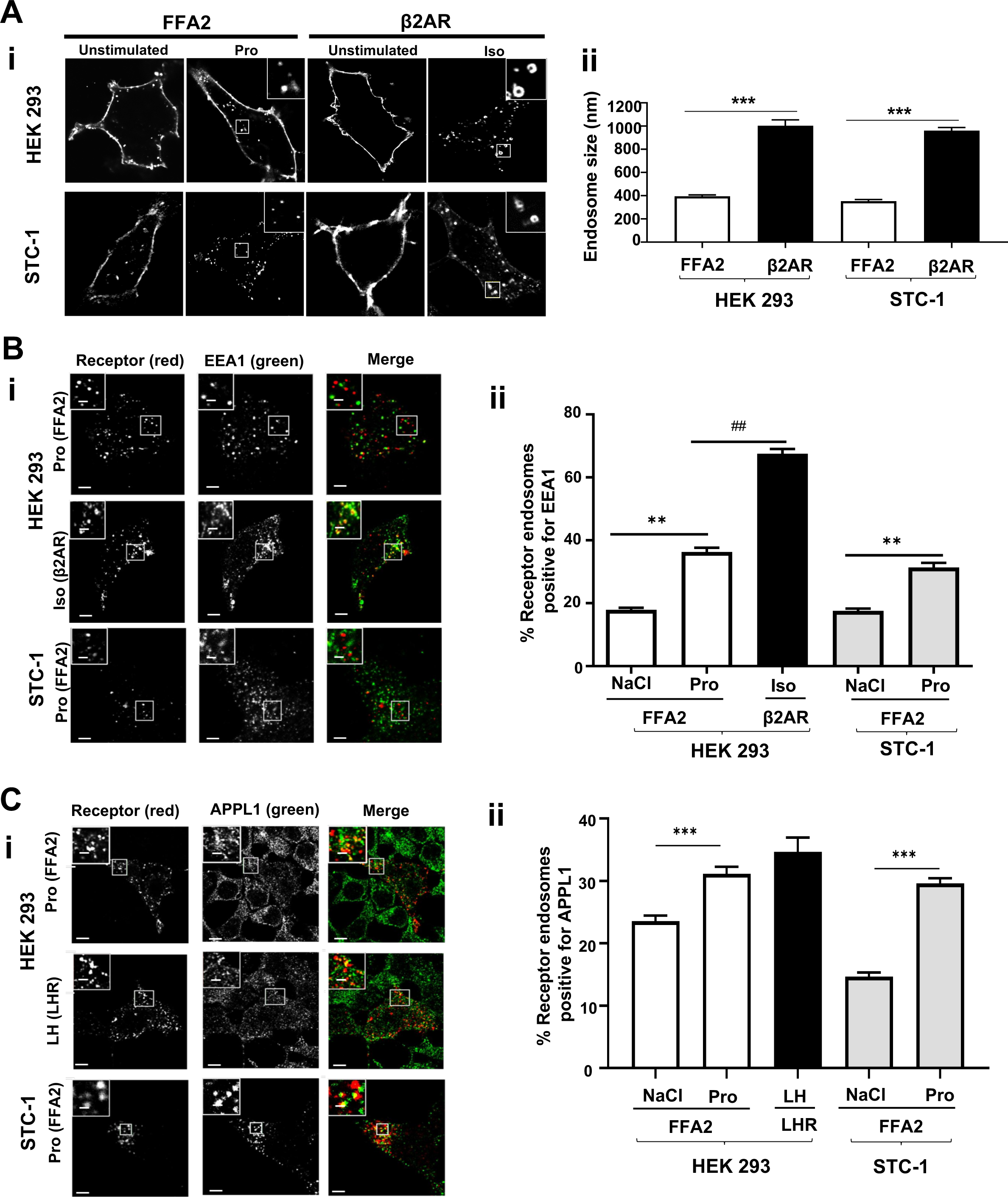
FFA2 internalizes to endosomes exhibiting properties of VEEs. (A) (i) Representative confocal microscopy images of HEK 293 cells expressing FLAG-FFA2 or FLAG-β2AR or STC-1 cells expressing FLAG-FFA2 or FLAG-β2AR imaged live with confocal microscopy before and after ligand treatment. FFA2 was stimulated with sodium propionate (Pro, 1mM), and β2AR with isoproterenol (Iso, 10 µM) for 20 min. Scale bars, 5 µm, scale bar in inset 1 µm. (ii) Bar graph showing diameter of FFA2 or β2AR in HEK 293 or STC-1 cells containing endosomes. Endosome diameter was assessed by measuring the diameter of 20 endosomes, n = 10 cells per condition, collected across 3 independent experiments. Two-sided Mann-Whitney U test, *** p < 0.001. (B) (i) Representative confocal microscopy images of fixed HEK 293 cells stably expressing FLAG-FFA2 or β2AR or STC-1 cells transiently expressing FLAG-FFA2 treated with ligand for 20 min prior to ‘stripping’ by PBS/EDTA (to remove surface bound FLAG antibody), fixation and stained with anti-EEA1 antibody. Scale bars, 5 µm, scale bar in inset 1 µm. (ii) Numbers of FFA2 or β2AR containing endosomes positive for EEA1 quantified from (i); 200 endosomes per condition, 10 cells quantified per condition. Data represent mean ± SEM, n=10 cells per condition, collected across 3 independent experiments. Two-sided Mann-Whitney U test, ** p < 0.01, *##* p <0.01. FFA2 colocalizes with APPL1. (i) Representative confocal microscopy images of fixed HEK 293 cells stably expressing FLAG-FFA2 or LHR or STC-1 cells transiently expressing FLAG-FFA2 treated with ligand (LH for LHR). Cells were treated as (B) except that cells were stained with anti-APPL1 antibody. Scale bars, 5 µm, scale bar in inset 1 µm. (ii) Numbers of FFA2 or LHR containing endosomes positive for APPL1 quantified from (i); 200 endosomes per condition, 10 cells quantified per condition. N = 10 cells per condition, collected across 3 independent experiments. Two-sided Mann-Whitney U test, *** p < 0.001. For confocal images, representative images are shown of ∼10 cells/experiment. Data represent mean ± SEM.

As FFA2 was primarily targeted to the VEE and propionate-induced FFA2 signaling requires internalization in enteroendocrine cells, we next investigated whether this compartment regulates FFA2 activity. We previously demonstrated that APPL1 is essential in rapid recycling of GPCRs targeted to this compartment back to the plasma membrane (Sposini et al., 2017). To determine a functional requirement of APPL1 on FFA2 trafficking, cellular levels of APPL1 were depleted via small interfering RNA (siRNA) in HEK 293 cells stably expressing FLAG-FFA2 (Figure 4A). We first examined the post-endocytic fate of FLAG-FFA2 when activated by propionate by confocal microscopy. APPL1 knockdown strongly impaired propionate-induced FFA2 recycling, in contrast, there was no effect in cells treated with NaCl (constitutive trafficking) as there was a complete return of the receptor back to the plasma membrane (Figure 4B). The role of APPL1 in rapid FFA2 recycling was quantitated via live-cell total internal reflection fluorescence microscopy (TIRFM) of an FFA2 tagged at the extracellular N-terminus with pH-sensitive GFP super-ecliptic pHluorin (SEP). SEP fluoresces in an extracellular neutral pH environment but is non-fluorescent when confined to the acidic lumen of endosomes, and therefore enables the detection of receptors upon insertion into the plasma membrane (Miesenbock et al., 1998; Yudowski et al., 2007). TIRFM imaging of SEP-tagged FFA2 (SEP-FFA2) established that FFA2 recycling events (identified as “puffs” of GFP fluorescence at the membrane) were transient and increased significantly within 5 minutes of propionate treatment, whereas treatment with NaCl exhibited a low rate of basal events (Figure S3A and B). The frequency of these events remained constant throughout the duration of the imaging period (30 minutes) (Figure S3C). These plasma membrane insertion events were not affected by pre-treatment of cells with cycloheximide, suggesting that such events are not due to *de novo* receptor biogenesis (Figure S3D). In cells depleted of APPL1, however, propionate-induced, but not constitutive, recycling of SEP-FFA2 was significantly impaired (Figure 4C).

**Figure 4.**
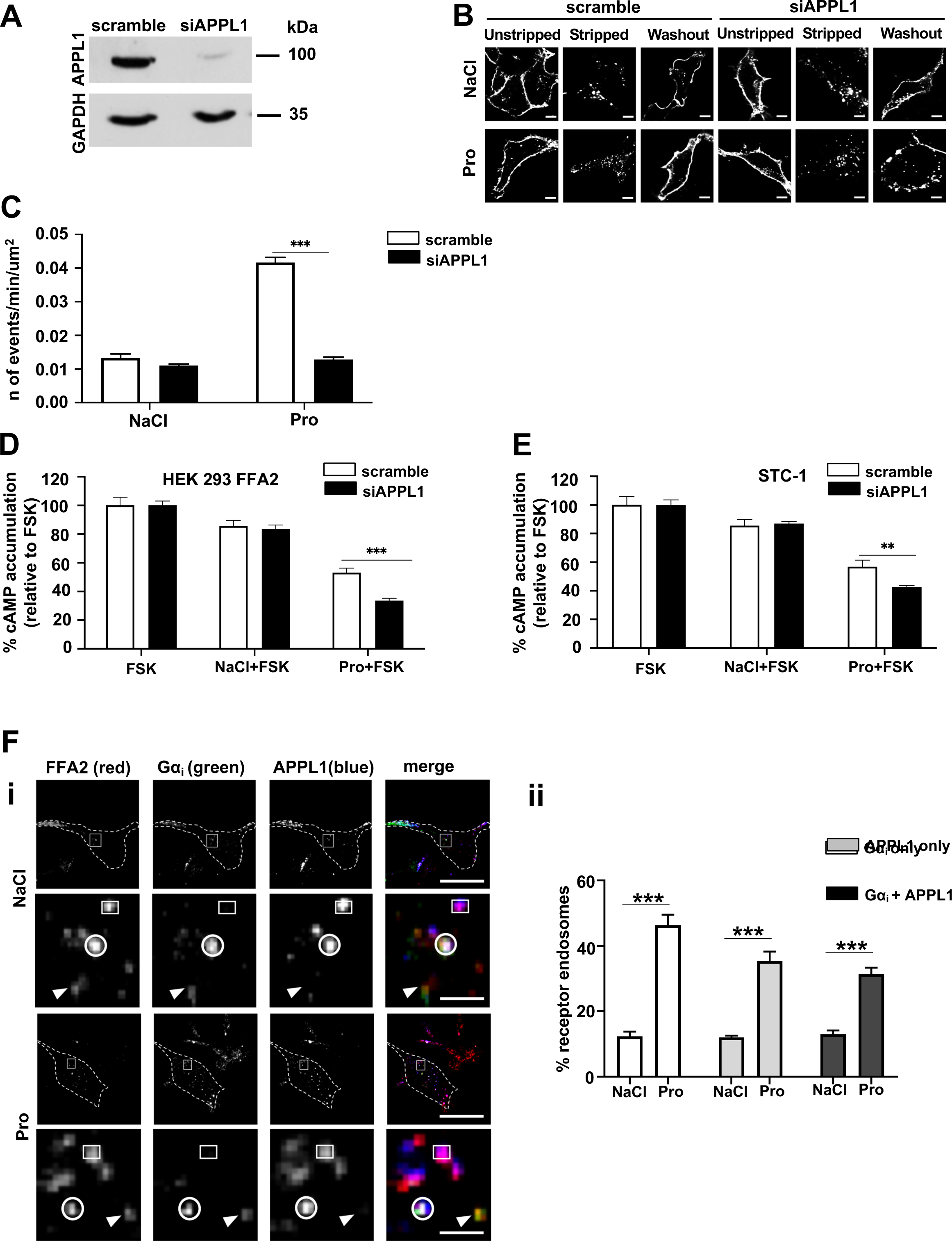
FFA2 trafficking and G protein signaling is regulated by APPL1. (A) Representative western blot of total cellular levels of APPL1 from cells transfected either with scramble or APPL1 siRNA. GAPDH was used as a loading control. (B) Representative confocal microscopy images of propionate-induced internalization and recycling following APPL1 siRNA-mediated knockdown. HEK 293 cells stably expressing FLAG-FFA2 were labelled with anti-FLAG antibody and then treated with NaCl (1 mM) or propionate (pro, 1 mM) for 20 min, then ‘stripped’ and incubated with ligand-free medium for 1h to allow receptor recycling. Scale bars, 5 µm, scale bar in inset 1 µm. (C) Recycling of HEK 293 cells stably expressing SEP-FFA2 was measured in real-time, via TIRFM, cells were transfected either with scramble or APPL1 siRNA and stimulated with NaCl (1 mM) or sodium propionate (Pro, 1 mM) for 5 min. n = 20 cells per condition, collected across 4 independent experiments. Two-sided Mann-Whitney U test, *** p < 0.001. (D) APPL1 negatively regulates propionate-mediated Gαi signaling. HEK 293 cells stably expressing FLAG-FFA2 (i) or STC-1 cells (ii) transfected with either scramble of APPL1 siRNA prior to pre-treatment of IBMX (0.5 mM, 5 min) and then stimulated with forskolin (FSK, 3 µM) or a combination of FSK and NaCl or stated SCFAs (1 mM, 5 min). Data are expressed as % change of FSK and NaCl treatment. n = 4 independent experiments. Two-sided Mann-Whitney U test, * p < 0.05; ** p < 0.01. (E) FFA2 colocalizes with Gα_i_ within APPL1 endosomes. Representative TIRFM images of HEK 293 cells stably expressing FLAG-FFA2 (red), Gαi (green), APPL1 (blue) in cells stimulated either with NaCl (1 mM) or sodium propionate (Pro, 1 mM) for 5 min (i). Dotted line marks cell boundary. The lower panel shows higher magnification image of the region of colocalization of the white box in the upper-panel images. Arrows indicate FFA2 endosomes positive for Gαi only; circle indicates FFA2 endosomes positive for Gαi and APPL1; squares indicated FFA2 endosome positive for APPL1 only. Scale bars of upper-panel images, 10 µm, scale bar of lower-panel images 3 µm. Quantification of FFA2 endosomes positive for either Gαi, APPL1 or Gαi and APPL1; n=12 cells per condition from (i) were quantified across 3 independent experiments (ii). Two-way ANOVA, Bonferroni multiple comparisons test, *** p < 0.001. Data represent mean ± SEM.

In addition to regulating the post-endocytic sorting of VEE targeted receptors, APPL1 also negatively regulates GPCR/Gαs signaling from this compartment (Sposini et al., 2017). As we have demonstrated that propionate-induced Gαi/o signaling requires receptor internalization, we next examined the potential role of the VEE in FFA2 signaling. Depletion of APPL1 resulted in a 2-fold increase in propionate-mediated inhibition of forskolin-stimulated cAMP (Figure 4D), suggesting that APPL1 can also negatively regulate heterotrimeric Gαi/o signaling in addition to Gαs signaling. Negative regulation of Gαi/o signaling by APPL1 was conserved in STC-1 cells as depletion of APPL1 levels also resulted in a significant enhancement of propionate-mediated inhibition of forskolin-induced cAMP (Figure S4 and Figure 4E).

The above data indicates that propionate-induced Gαi/o signaling may occur from VEEs, and regulated by APPL1 endosomes, therefore we next determined whether FFA2 colocalizes with Gαi in APPL1-positive endosomes. HEK 293 cells expressing FLAG-FFA2 and Gαi-venus were imaged via TIRFM as VEEs are prevalent in the peripheral juxtamembrane region of cells (Sposini et al., 2017). TIRFM analysis revealed that FFA2 and Gαi-venus positive endosomes are heterogeneous and characterized by FFA2-Gαi endosomes with and without APPL1. In addition, FFA2 endosomes were also positive for APPL1 where no Gαi-was present (Figure 4F). Overall, these data demonstrate that APPL1 is essential for propionate-induced FFA2 trafficking from the VEE and regulation of propionate-mediated Gαi/o signaling.

### Endosomal Gαi/o signaling regulates propionate-induced GLP-1 release

As FFA2 internalization requires propionate-induced Gαi/o signaling, we next assessed whether Gαi/o endosomal signaling regulates GLP-1 secretion. First, we examined the involvement of Gαi/o versus Gαq/11 signaling in mediating propionate-induced GLP-1 release, using Gαi/o or Gαq/11 inhibitors at concentrations that we demonstrated could inhibit receptor signaling in enteroendocrine cells (Figure 1A-B and Figure S1B). STC-1 cells or colonic crypts with pretreated with Ptx impaired propionate-induced GLP-1 release (Figure 5A-B). In contrast, pretreatment of STC-1 cells with the Gαq/11 inhibitor, YM-254890, had no significant effect on propionate-mediated GLP-1 release (Figure 5C). In colonic crypts however, propionate-induced GLP-1 release in the presence of YM-25480 was impaired (Figure 5D). As we did not observe propionate-induced Gαq/11 signaling in colonic crypts (Figure 1F-I), we determined if propionate-mediated FFA2 Gαi/o signaling is altered in the presence of YM-25480 in these primary cultures. In the presence of YM-25480, propionate-mediated inhibition of forskolin-induced cAMP was partially, but significantly impaired compared to control treated cells (Figure S5A). This was only specific to propionate-mediated FFA2 signaling, as YM-25480 had no effect on propionate-mediated signaling from FFA3, a receptor coupled to only Gαi/o (Figure S5B). Together these data suggest that in STC-1 cells and colonic crypts, propionate induces GLP-1 secretion via a Gαi/o-dependent mechanism.

**Figure 5.**
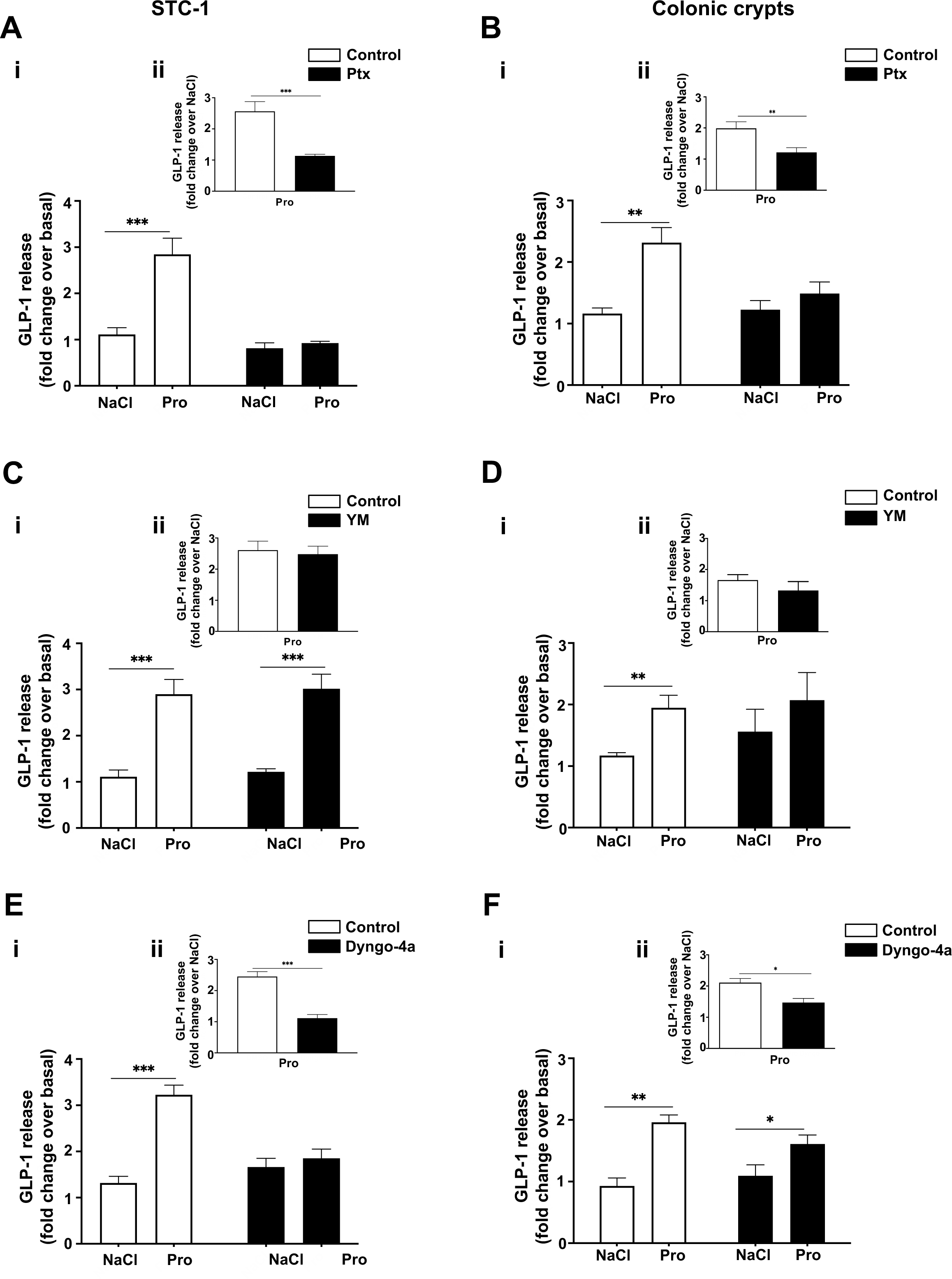
Endosomal Gαi/o signaling regulates propionate-mediated GLP-1 release. Stimulation of GLP-1 release from STC-1 cells (A) or colonic crypts (B) in the presence of Ptx. STC-1 cells or colonic crypts were pre-treated with either vehicle or Ptx (200 ng/mL, 20 h) prior to stimulation with either NaCl (1 mM) or sodium propionate (Pro, 1 mM) for 2 h and 1 h for colonic crypts. For STC-1 cells, n=4 independent experiments. For crypts, n=8 independent experiments. Two-sided Mann-Whitney U test, *** p < 0.001. Stimulation of GLP-1 release from STC-1 cells (C) or colonic crypts (D) in the presence of Gαq/11 inhibitor, YM-254890. STC-1 cells and colonic crypts were pre-treated with either DMSO or YM-254890 (YM, 10 µM, 5 min) and then treated as in (A and B). For STC-1 cells, n = 4 independent experiments. For crypts, n=3 independent experiments. Two-sided Mann-Whitney U test, * p < 0.05, ** p <0.01. Stimulation of GLP-1 release from STC-1 cells (D) or colonic crypts (E) in the presence of Dygno-4a. STC-1 cells or colonic crypts were pre-treated with either DMSO or Dyngo-4a (50 µM, 45 min for STC-1 cells and 100 µM, 45 min for colonic crypts), following pre-treatment, Dyngo-4a was co-incubated with ligands for an additional 5 min and then removed. Cells and crypts were treated as (A and B). For STC-1 cells, n = 5 independent experiments. For crypts, n = 3 independent experiments. Two-sided Mann-Whitney U test, ** p < 0.01, *** p <0.001. Insets show propionate-induced GLP-1 release normalized to NaCl GLP-1 release. ** p < 0.01, *** p <0.001. GLP-1 secretion of media and cells were detected via RIA and was expressed as fold change of total GLP-1 and normalized to NaCl secretion within the same experiment. Data represent mean ± SEM.

The requirement of Gαi/o signaling for propionate-mediated GLP-1 release, and the critical role of propionate-driven FFA2 internalization for G protein signaling suggests a role for propionate-induced receptor internalization. To test this hypothesis, receptor endocytosis in STC-1 cells was blocked by Dyngo-4a treatment (Figure 2G). In cells treated with Dyngo-4a, propionate exhibited a marked reduction in GLP-1 release compared to control treated cells (Figure 5E). This inhibition by Dyngo-4a was not a result of an overall decreased capacity for these cells to secrete hormone as forskolin-induced GLP-1 release was not affected by inhibition of dynamin GTPase activity (Figure S6A). In colonic crypts, pretreatment with Dyngo-4a, impaired propionate-induced GLP-1 release but not to the same degree as observed in STC-1 cells (Figure 5F). As we have observed an essential dependence of propionate-driven FFA2 internalization for not only G protein signaling in STC-1 cells (Figure 2H) but also for GLP-1 release, we hypothesized that the lack of modulation of propionate-induced GLP-1 release in the presence of Dyngo-4a in colonic crypts may be due to more technical limitations of Dyngo-4a in primary tissue compared to a monolayer of cells. To assess this, we determined the ability of propionate to inhibit forskolin-induced cAMP in the presence of Dyngo-4a in colonic crypts. In the presence of Dyngo-4a, propionate-mediated inhibition of forskolin-induced cAMP was significantly, but only partially impaired compared to control treated cells (Figure S6B), in contrast to the full inhibition of Gαi/o signaling by Dyngo-4a observed in STC-1 cells (Figure 2H). Thus, the level of propionate-dependent Gαi/o signal inhibition by Dyngo-4a correlates with its ability to inhibit propionate-driven gut hormone secretion. Overall, this suggests that endosomal Gαi/o signaling mediates propionate-induced GLP-1 release.

### Propionate-induced endosomal signaling regulates GLP-1 release via activation of p38

Increases in intracellular cAMP is an established driver of gut hormone release, yet our data indicates propionate-induces gut hormone release in a Gαi/o-dependent manner, a pathway that decreases cAMP levels. Therefore, we hypothesized that the mechanism mediating endosomal Gαi/o-dependent GLP-1 release is potentially via distinct downstream pathways activated by Gαi/o, rather than its actions on its effector enzyme adenylate cyclase. Thus, we determined which propionate-mediated signaling pathways downstream of G protein signaling are also spatially regulated. A phosphokinase array was employed in STC-1 cells to identify propionate-induced signaling pathways dependent on receptor internalization. STC-1 cells were pretreated with Dyngo-4a and stimulated with propionate for 5 or 30 minutes. The array revealed that 16 of the 43 kinases within the array were phosphorylated after 5 or 30 min of propionate treatment. However, only p38α, EGF-R, MSK1/2 and Hck showed reduced phosphorylation when internalization was inhibited (Figure 6A). Of these kinases, p38α was selected for further analysis as this kinase and MSK1/2 are part of the same signal cascade. Furthermore, p38α is known to be activated at endosomes by other GPCRs (Grimsey et al., 2015). We then asked if propionate-induced p38 activation was Gαi/o-mediated. To test this, propionate-induced p38 activation was assessed in STC-1 pretreated with Ptx via Western blot. Pretreatment of Ptx significantly impaired propionate-induced p38 signalling (Figure 6B).

**Figure 6.**
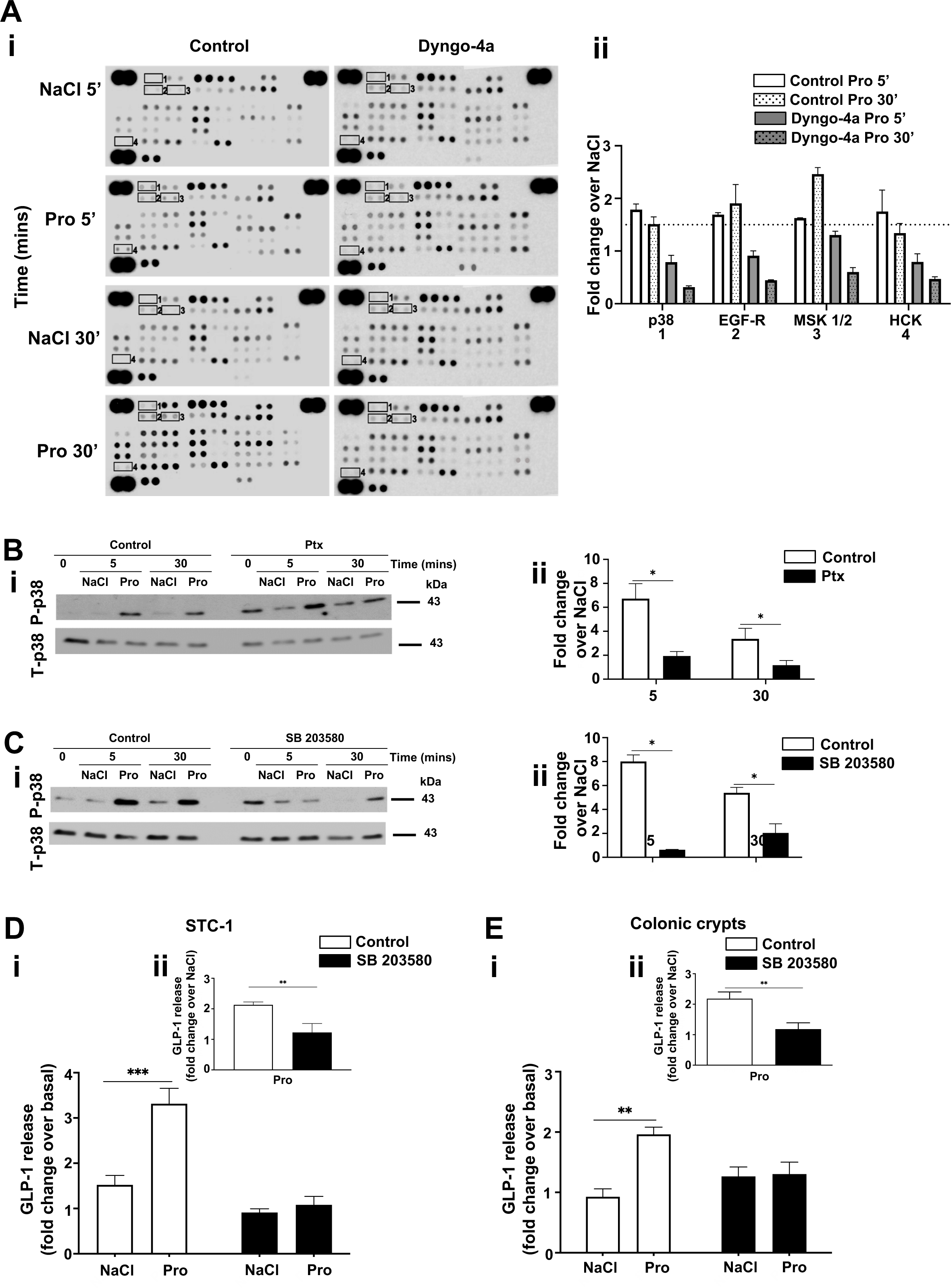
Endosomal signaling of FFA2 regulates GLP-1 release via activation of p38. (A) STC-1 cells were pre-treated with DMSO (vehicle) or Dyngo-4a (50 μM, 45 mins) prior to stimulation with NaCl (1 mM) or propionate (Pro, 1 mM) for 5 or 30 min. Lysates were incubated with membranes spotted for 43 different phosphokinases (R&D systems). (Ai) Membranes highlighting location of kinase phospho-antibodies spotted onto the array. Signals of relevant kinases in response to Dyngo-4a effects are indicated by numbers. (Aii) Fold changes over NaCl in levels of phosphorylation that decreased in presence of Dyngo-4a. Data represent mean ± SEM of fold change values. (B-C) Representative Western blot demonstrating phosphorylated p38 (P-p38) and total p38 (T-p38) of lysates from STC-1 cells with p38 inhibitor, SB 203580 (A), Dyngo-4a (B), Ptx (C). STC-1 cells were pre-treated with DMSO (vehicle) or SB 203580 (5 μM, 10 min), Dygno-4a (50 μM, 45 min), Ptx (200 ng/mL, 20 hours) prior to stimulation of NaCl (1 mM) or propionate (Pro, 1 mM) at the indicated time points. Cell lysates were then collected for Western blot analysis and probed for P-p38. Membranes were then stripped and re-probed with t-p38 which was used as a loading control (i). Densitometry and fold change analysis of P-p38 normalized to total-p38 of lysates pre-treated with control, Ptx, or SB 203580. Fold change of densitometry analysis of P-p38 levels normalized to NaCl of control or inhibitor at each time point stimulation with total-p38 (ii). Stimulation of GLP-1 release from STC-1 cells (D) or colonic crypts (E) in the presence of SB 203580. Both were pre-treated either with DMSO or SB 203580 (5 μM, 10 min), prior to stimulation with either NaCl (1 mM) or sodium propionate (Pro, 1 mM) for 2 h for STC-1 cells and 1 h for colonic crypts. For STC-1 cells, n = 3 independent experiments. For crypts, n=3 independent experiments. Two-sided Mann-Whitney U test, *** p < 0.001. Insets show propionate-induced GLP-1 release normalized to NaCl-induced GLP-1 release. ** p < 0.01, *** p <0.001. GLP-1 secretion of media and cells detected via RIA and was expressed as fold change of total GLP-1 and normalized to NaCl secretion within the same experiment. Data represents mean ± SEM.

Since propionate-induced activation of p38 involves receptor internalization and Gαi/o signaling, its role in propionate-induced GLP-1 release was assessed. A widely used selective p38 inhibitor SB 203580, which inhibits the catalytic activity of p38-α and - β isoforms without inhibiting p38 phosphorylation mediated by upstream kinases (Ge et al., 2002) was employed and significantly impaired propionate-induced activation of p38 (Figure 6C). In STC-1 cells and colonic crypts, SB 203580 pretreatment significantly impaired propionate’s ability to induce GLP-1 secretion (Figure 6D-E) suggesting that propionate-induced endosomal Gαi signaling regulates GLP-1 release via a p38-dependent mechanism.

## DISCUSSION

The ability of propionate to stimulate the release of anorectic gut hormones via the GPCR FFA2 represents a key physiological function of high interest due to its demonstrated health benefits (Chambers et al., 2015; Chambers et al., 2019). However, despite our increasing knowledge of the complexity of GPCR signaling networks in other cell systems, the underlying mechanisms regulating gut hormone release by propionate/FFA2 are poorly understood. In this study we demonstrate that signaling and downstream functions of FFA2, in response to propionate is specified through tight control of receptor location.

In the gut, the current view is that GPCRs coupled to either Gαs-cAMP or Gαq/11-calcium pathways mediate anorectic gut hormone release (Hauge et al., 2017; Tian and Jin, 2016). From a receptor perspective, however, it is well known that many GPCRs are pleiotropically coupled, either directly or via receptor crosstalk, and where additional mechanisms, such as intracellular receptor signaling, enable diversity in cell functions from the same G protein and second messenger system. Furthermore, different GPCR ligands (endogenous and synthetic) can elicit distinct conformational states, and thus the potential to induce bias signal activity from the same receptor. In regard to FFA2 activity, which has been characterized previously as a dually coupled GPCR in studies primarily in heterologous cells (Brown et al., 2003; Le Poul et al., 2003), to date there have been no studies demonstrating its pleiotropic coupling to both Gαi/o and Gαq/11 in the gut at the level of second messenger signaling. Thus, to delineate the mechanisms of propionate-induced GLP-1 release from enteroendocrine cells, we first profiled the second messenger signaling activated by this SCFA in our intestinal models. While propionate robustly signals via Gαi/o in a FFA2-dependent manner, it was unable to induce Gαq/11 signaling both in colonic crypts and STC-1 cells. This is in contrast to prior studies reporting a propionate-dependent calcium response in colonic cultures expressing Venus fluorescent protein in enteroendocrine L cells (Tolhurst et al., 2012). The reasons for this disparity are unclear, but could relate to either the mouse model harboring Venus protein, longer culture times employed to create dispersed colonic cultures to measure calcium signaling, as opposed to the intact colonic crypts used in this study, and/or reflect a Gαi/o-mediated response as Gαi/o-coupled GPCRs are known to modulate calcium responses, including influx of extracellular calcium (Tang et al., 2015; Alkhatib et al., 1997). To our knowledge this is also the first demonstration that previously characterized synthetic orthosteric and allosteric FFA2 selective ligands activate Gαq/11 signaling in the colon. Although propionate may also activate FFA3, a Gαi/o-coupled receptor known to also be expressed in the colon, it was demonstrated in this study that colonic crypts from FFA2 KO animals are unable to activate SCFA-dependent Gαi/o signaling, and that these animals do not exhibit altered Ffar3 levels. This supports prior published work from us and others demonstrating SCFA-mediated GLP-1 release requires FFA2 (Tolhurst et al., 2012; Psichas et al., 2015). FFA2 is known to be a dually coupled receptor in HEK 293 or CHO cells (Le Poul et al., 2003), and indeed our data in HEK 293 cells expressing FFA2 is consistent with these reports, whereby propionate also activates Gαq/11 signaling. This potential system-dependent bias exhibited by FFA2 and propionate is intriguing given FFA2 can activate Gαq/11 signaling in enteroendocrine cells when stimulated with synthetic ligands. One potential mechanism for the distinct propionate/FFA2 signal profiles between enteroendocrine cells and heterologous cells is crosstalk of FFA2 with another GPCR such as FFA3. However, it has recently been demonstrated that FFA2-Gαq/11 signaling is not decreased, but enhanced, via associations with FFA3 (Ang et al., 2018).

Propionate’s inability to induce Gαq/11-mediated intracellular calcium mobilization is indeed paradoxical to what is known about the signaling requirements of anorectic gut hormone secretion (Spreckley and Murphy, 2015). However, exocytosis is known to occur via either calcium-dependent and -independent pathways (Sato et al., 1998; Komatsu et al., 1995) and also involving Gαi (Aridor et al., 1993). Although we observed that propionate-induced GLP-1 release was impaired in the presence of Ptx in STC-1 cells and colonic crypts, this is inconsistent with previous reports that did not find propionate-induced GLP-1 release to be modulated by Ptx (Tolhurst et al., 2012; Bolognini et al., 2016), but is impaired by the Gαq/11 inhibitor, FR900359 (Bolognini et al., 2016). However, confirmation of Ptx-dependent inhibition of propionate-mediated Gαi/o signaling at the second messenger level in STC-1 cells or crypts was not reported in these prior studies. Although FR900359 has been reported to also inhibit Gβγ-mediated signaling from Gαi/o-coupled receptors (Gao and Jacobson, 2016), we observed that inhibition of Gαq/11 activation partially inhibited propionate-FFA2 Gαi/o signaling, suggesting that an active Gαq/11 is integrated with FFA2 Gαi/o signaling. Such crosstalk may be analogous to the findings that arrestin-mediated signaling of GPCRs requires an active G protein state perhaps even in the absence of second messenger responses (Grundmann et al., 2018).

Our results demonstrating propionate-mediated GLP-1 release via Gαi/o suggest that mechanisms regulating propionate-induced release of GLP-1 may be more complex and not via Gαi/o-mediated decreases in cAMP levels per se, a second messenger that induces gut hormone release (Hauge et al., 2017). One mechanism that can diversify downstream cellular functions from a common upstream pathway is via spatial control of signaling. Indeed, agonist-induced FFA2 internalization differentially regulated Gαi and Gαq/11 signaling, demonstrating at least in HEK 293 cells that FFA2/Gαi signaling was endosomal and Gαq/11 signaling occurred from the plasma membrane. Spatial discrimination in GPCR/G protein signaling has been observed with the pleiotropically-coupled calcium-sensing receptor (Gorvin et al., 2018). The specific requirement for receptor internalization in driving FFA2-mediated Gαi/o signaling was conserved in enteroendocrine cells, whereby propionate-mediated Gαi/o signaling, and GLP-1 release required internalization, providing a novel mechanism underlying propionate’s downstream functions in the gut. We also identified that FFA2 primarily traffics to the VEE, an endosomal compartment we have previously shown is critical for sorting and endosomal signaling for a subset of GPCRs (Jean-Alphonse et al., 2014; Sposini et al., 2017). For GPCRs that are targeted to the VEE, APPL1 has been demonstrated to be crucial for both receptor recycling and negative regulation of Gαs signaling. We demonstrate that rapid ligand-induced recycling of FFA2 is also APPL1-dependent and negatively regulates FFA2-endosomal Gαi/o signaling in HEK 293 and enteroendocrine cells, indicating that the APPL1/VEE compartment can negatively regulate distinct G protein pathways, in addition to Gαs-coupled GPCRs (Sposini et al., 2017).

Given the requirement for active endosomal Gαi/o in mediating propionate-induced gut hormone release, we hypothesized that additional endosomal Gαi/o-activated pathways were important in gut hormone secretion. We identified that phosphorylation of a small subset of downstream kinases required FFA2 internalization when activated by propionate, which supports a role for endomembrane signaling in providing a signal platform to activate unique signaling substrates from the plasma membrane, or indeed other intracellular compartments (Eichel and von Zastrow, 2018; Hanyaloglu, 2018). We focused on p38 as kinases of the same pathway, MSK1/2, were also identified in the array, and p38 is known to be activated endosomally by other GPCRs (Grimsey et al., 2015). Propionate has also previously been shown to activate p38 in many cellular systems and have a role in regulating inflammatory responses (Rutting et al., 2019; Yonezawa et al., 2007; Ang et al., 2018). For enteroendocrine cells and colonic crypts, we identify a key role of p38 in regulating propionate-induced GLP-1 release. Interestingly, p38 is also involved in regulating GLP-1 secretion induced by meat hydrolysate and essential amino acid- and low molecular weight chitosan (Reimer, 2006; Liu et al., 2013). More recently, propionate-induced GLP-1 release was also found to be regulated by p38 in chicken intestinal epithelial cells, (Zhang et al., 2019), suggesting a conserved role of this kinase in anorectic gut hormone secretion induced by distinct metabolites.

Together, these findings strongly support a model whereby the unique health benefits of propionate to regulate appetite requires tightly controlled integration of membrane trafficking and endosomal signaling. Such a model offers a future platform to evaluate specific populations, pathophysiological alterations and the long-term health potential of elevated colonic propionate. Furthermore, it may represent a broader mechanism employed by intestinal metabolites, which activate multiple GPCRs within the gut, to diversify its functions *in vivo*.

## AUTHOR CONTRIBUTIONS

N.C performed all signaling and trafficking experiments and GLP-1 studies in STC-1 cells. N. G-A and Y.M prepared colonic crypt cultures. N. G-A carried out GLP-1 experiments in colonic crypts and Y.M genotyped and assessed Ffar3 levels in crypts. M.S. synthesized and characterized Cmp1 under supervision of E.W.T. A.I contributed the β-arrestin1/2 KO cell line. N.C, N. G-A, E.W.T., G.F and A.C.H designed research, and with N.C, N. G-A, A.I, Y.M E.W.T and M.S analyzed data and wrote the paper. All authors critically read and approved the final manuscript.

## ACKNOWLEDGEMENTS

We would like to thank Drs. Andreas Bruckbauer and Stephen Rothery at the Facility for Imaging of Light Microscopy at Imperial College London for technical support with TIRFM and Dr. Paul Bech and Prof. Kevin Murphy (Imperial College London) for assistance with RIAs. FLAG-FFA3 plasmid was provided by Ms. Tilly Shackley (Imperial College London). This work was supported by grants from the Biotechnology and Biological Sciences Research Council to G.F, A.C.H and E.T.W (BB/N016947/1) and to A.H and E.T.W (BB/S001565/1). A.I. was funded by the PRIME (JP18gm5910013) and the LEAP (JP18gm0010004) from the Japan Agency for Medical Research and Development (AMED); JSPS KAKENHI grant (17K08264) from the Japan Society for the Promotion of Science

## MATERIALS AND METHODS

### Animals

C57BL/6J mice purchased from Charles River were used to prepare mouse colonic crypts. FFA2 global knockout (FFA2 ^-/-^) mice were generated by Deltagen. FFA2 knockout was achieved by homologous recombination that replaces 55bp of FFA2 exon 1 with a cassette containing the neomycin resistance and β-galactosidase genes, resulting in a frameshift mutation (Maslowski et al., 2009). Animals were cared for in accordance with British Home Office under UK Animal (Scientific Procedures) Act 1986 (Project License 00/6474).

### Mouse colonic crypt culture preparation

Colons of male wildtype (WT) or FFA2 ^-/-^ C57BL6 mice (8-12 weeks of age) were removed, cleaned and placed into ice-cold L-15 (Leibowitz) medium. The intestinal tissue was thoroughly cleaned with L-15 medium and digested with 0.4 mg ml-1 collagenase in high-glucose DMEM at 37°C, as described previously (Psichas et al., 2015).). The digestion process was repeated 4 times and resulting cell suspensions were centrifuged (5 min, 300 g). The pellets were resuspended in DMEM (supplemented with 10% fetal calf serum and 1% antibiotics, 100 U ml-1 penicillin and 0.1 mg ml-1 streptomycin). Combined cell suspensions were filtered through a nylon mesh (pore size ∼250 μm) and plated onto appropriate culture plates, 2% Matrigel-coated plates. The plates were incubated overnight at 37°C in an atmosphere of 95% O2 and 5% CO_2_.

### Colonic crypt FFA3 mRNA expression levels

Total RNA was extracted from WT and FFA2^-/-^ (age-matched) plated colonic crypts using PureLink® RNA Mink Kit (Invitrogen) and DNase treated using on-column PureLink® DNase Treatment (Invitrogen). DNase-treated total RNA was reversed transcribed to a single-stranded cDNA using the high-capacity cDNA Reverse Transcription kit (Applied Biosystems). Quantitative reverse transcriptase PCR (qPCR) was carried out by QuantStudio® 12 K Flex Real-Time PCR System (Life Technologies) using TaqMan Gene Expression Assay (Applied Biosystems) with FFAR3 hydrolysis probe (Mm02621638_1, Applied Biosystems) and 18S as the reference gene (Eukaryotic 18S rRNA Endogenous Control, Applied Biosystems). The qPCR data are presented as relative expression levels calculated by ΔΔCt (where ΔCt is determined by the difference cycles threshold of the target gene and the reference gene).

### Synthesis of Compound 1

All solvents and reagents were purchased from Sigma-Aldrich, Alfa Aesar unless otherwise stated, and used without further purification. Moisture sensitive reactions were performed in oven dried flasks, under a nitrogen atmosphere. Anhydrous solvents were dispensed using Pure Solv™ solvent drying towers (Innovative Technology Inc.) Analytical thin layer chromatography was carried out using Merck Si_60,_ F_254_ chromatography sheets. Spots were visualised by UV light or through use of an appropriate stain (ninhydrin or potassium permanganate). Flash column chromatography was run on a Biotage Isolera™ One flash purification system using a wet-loading Biotage SNAP cartridge. Mass spectra were acquired by the Imperial Mass Spectrometry service with *m/z* values reported in Daltons. ^1^H spectra were recorded on a Bruker Av-400 (400 Hz) instrument at RT. Chemical shifts are expressed in parts per million δ relative to residual solvent as an internal reference. The multiplicity if each signal is indicated by: s = singlet; broad s= broad singlet; d = doublet; t = triplet; m= multiplet. Coupling constants (*J*), calculated using MestReNova^©^ NMR software, are quoted in Hz and recorded to the nearest 0.1 Hz.

### 1 *N*-cyclopropyl-*N*’-benzoylthiourea

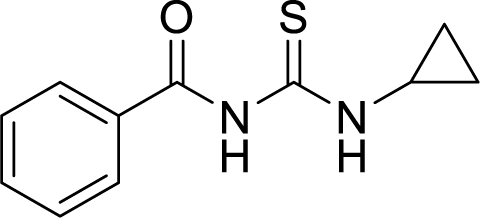

Benzoyl isothiocyanate (820 μL, 6.13 mmol, 1.0 eq.) was dissolved in CH_2_Cl_2_ (25 mL) at 0 °C, followed by a dropwise addition of cyclopropylamine (425 μL, 6.13 mmol, 1.0 eq.). The solution was then warmed up to RT and allowed to stir for 17h. The crude mixture was concentrated *in vacuo*, yielding benzoylthiourea as a yellow solid (1348 mg, quant.), which was used in the next step without further purification. ^1^H NMR (400 MHz, CDCl_3_): δ 10.93 (1H, broad t), 9.17 (1H, s), 7.83 (2H, dd, *J* = 8.4, 1.4 Hz), 7.59 (1H, t, *J* = 7.4 Hz), 7.48 (2H, t, *J* = 7.9 Hz), 3.23-3.17 (1H, m), 0.93-0.89 (2H, m), 0.80-0.77 (2H, m). The compound has been characterized in the literature, data in agreement (Olken and Marletta, 1992).

### 2 Cyclopropylthiourea

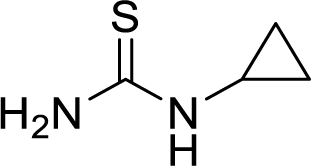

Benzoylthiourea **1** (900 mg, 4.09 mmol, 1 eq.) was dissolved in a solution of 5% (w/v) NaOH (20 mL) and heated to 80 °C. The solution was stirred for 3h and then cooled to RT in an ice/water bath. The reaction mixture was titrated to pH 8.0 with HCl_conc_. The crude was extracted with EtOAc (4 × 15 mL). The organic fractions were combined, dried over anhydrous MgSO_4_, filtered and concentrated *in vacuo*. The crude was redissolved in CH_2_Cl_2_ (10 mL) and precipitated with a dropwise addition of Et_2_O to afford a thiourea **2** as a white-off solid (200 mg, 42%), which was used in the next step without further purification. ^1^H NMR (400 MHz, DMSO-d_6_): δ 2.35 (1H, broad s), 0.67-0.63 (2H, m), 0.49-0.44 (2H, m). The compound has been characterized in the literature, data in agreement (Olken and Marletta, 1992).

### 3 2-Bromo-1-(2,5-dichlorophenyl)ethanone

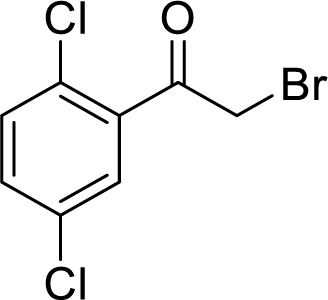

Dichloroacetophenone (230 μL, 1.59 mmol, 1.0 eq.) was dissolved in anhydrous MeCN (8 mL) and cooled to 0 °C under nitrogen, then NBS (312 mg, 1.75 mmol, 1.1. eq.) was added, followed by a dropwise addition of TMS·OTf (14 μL, 0.08 mmol, 0.05 eq.). The solution was warmed up to RT and allowed to stir for 17h under nitrogen. The reaction mixture was concentrated *in vacuo* and purified by column chromatography (1 to 5% EtOAc in Hexane over 10 CV), which afforded bromide as a white-off thick oil (235 mg, 55%, 80% pure by NMR). ^1^H NMR (400 MHz, CDCl_3_): δ 7.54 (1H, d, *J* = 2.4 Hz), 7.40 (1H, d, *J* = 2.3 Hz), 7.39 (1H, s), 4.49 (2H, s). The compound has been characterized in the literature, data in agreement (Roman et al., 2010)

### *4 N*-cyclopropyl-4-(2,5-dichlorophenyl)thiazol-2-amine

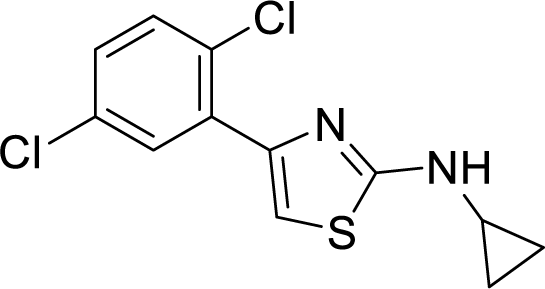

Cyclopropylthiourea **2** (50 mg, 0.43 mmol, 1.0 eq.) was dissolved in ethanol (2 mL), followed by addition of bromide **3** (80% pure, 138 mg, 0.52 mmol, 1.2 eq.) pre-dissolved in ethanol (1 mL). The solution was allowed to stir for 3h at RT and then concentrated *in vacuo*. The residue was dissolved in CH_2_Cl_2_ (5 mL), washed with saturated NaHCO_3_ (4 mL), brine (4 mL). Organic layer was dried over MgSO_4_, filtered and concentrated *in vacuo* to give a thick yellow oil. Column chromatography (1 to 10% EtOAc in Hexane over 10 CV) afforded amine **4** as an off-white thick oil (92 mg, 75%). R_f_ 0.57 (Hex:EtOAc = 3:1). ^1^H NMR (400 MHz, CDCl_3_): δ 7.86 (1H, d, *J* = 2.6 Hz), 7.37 (1H, d, *J* = 8.4 Hz), 7.19 (1H, dd, *J* = 8.3, 2.8 Hz), 7.13 (1H, broad s), 7.08 (1H, s), 2.60-2.54 (1H, m), 0.69-0.64 (2H, m), 0.56-0.52 (2H, m). The compound has been characterized in the literature, data in agreement (Hoveyda et al., 2018).

### *5 tert*-Butyl (R)-3-benzyl-4-(cyclopropyl(4-(2,5-dichlorophenyl)thiazol-2-yl)amino)-4-oxobutanoate

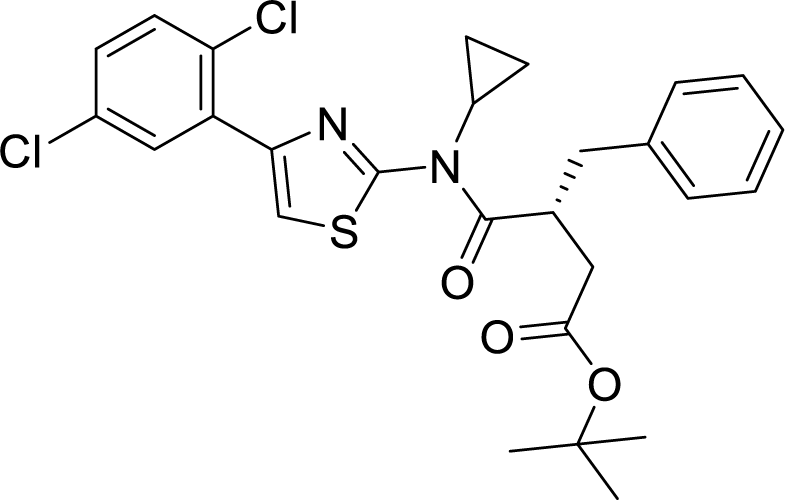

In a dry microwave vial (R)-2-benzyl-4-(tert-butoxy)-4-oxobutanoic acid (30 mg, 0.114 mmol, 1.3. eq.) was dissolved in anhydrous CH_2_Cl_2_ (1 mL) under nitrogen. Fluoro-*N,N,N’,N’*-bis(tetramethylene)formamidinium hexafluorophosphate (BTFFH) (41 mg, 0.131 mmol, 1.5 eq.) was then added, followed by anhydrous i-Pr_2_NEt (68 μL, 0.391 mmol, 4.5. eq.). The solution was allowed to stir for 30 min at RT under nitrogen, followed by addition of amine **4** (25 mg, 0.088 mmol, 1.0 eq.). The vial was then sealed, heated to 80 °C in an oil bath and allowed to stir for 18h. The yellow reaction mixture was cooled down to RT, further diluted with CH_2_Cl_2_ (5 mL), quenched with saturated NH_4_CL (4 mL), washed with H_2_O (4 mL) and brine (4 mL). The organic layer was dried over anhydrous MgSO_4_, filtered and concentrated *in vacuo* to give a dark-yellow thick oil. Column chromatography (1 to 10% EtOAc in Hexane over 10 CV) afforded amide **5** as a light-yellow thick oil (37 mg, 78%). R_f_ 0.65 (Hex:EtOAc = 3:1). ^1^H NMR (400 MHz, CDCl_3_): δ 7.99 (1H, d, *J* = 2.8 Hz), 7.62 (1H, s), 7.39 (1H, d, *J* = 8.6 Hz), 7.30-7.15 (6H, m), 4.23-4.13 (1H, m), 3.08 (1H, dd, *J* = 13.4, 6.7 Hz), 2.96-2.82 (2H, m), 2.73 (1H, dd, *J* = 13.4, 8.2 Hz), 2.44 (1H, dd, *J* = 16.4, 4.9 Hz), 1.39 (9H, s), 1.30-1.20 (3H, m), 0.85-0.78 (1H, m). LRMS (ES^+^): 531 ([^35^Cl^35^ClM+H]^+^, 100%), 533 ([^35^Cl^37^ClM+H]^+^, 75%), 535 ([^37^Cl^37^ClM+H]^+^, 20%). The compound has been characterized in the literature, data in agreement (Hoveyda et al., 2018).

### (R)-3-benzyl-4-(cyclopropyl(4-(2,5-dichlorophenyl)thiazol-2-yl)amino)-4-oxobutanoic acid (Cmp1)

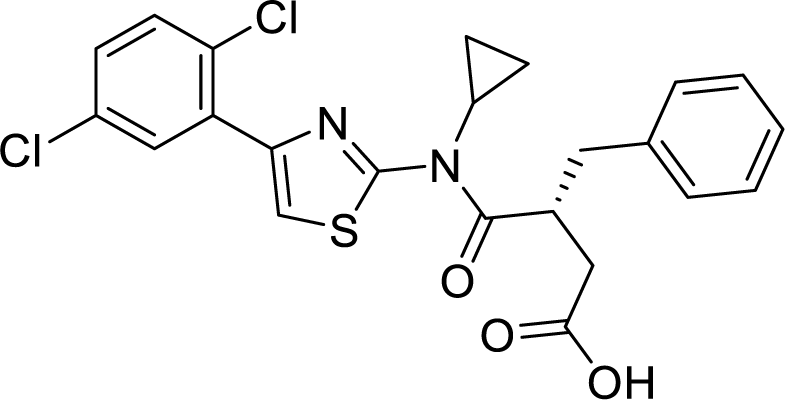

Amide **5** (10 mg, 0.019 mmol, 1.0 eq.) was dissolved in anhydrous CH_2_Cl_2_ (180 mL) under nitrogen, followed by addition of TFA_conc_ (40 mL, 20% (v/v)). The solution was allowed to stir for 4h at RT. The reaction mixture was concentrated *in vacuo*, redissolved in CH_2_Cl_2_ (3 mL) and quenched with saturated NaHCO_3_ (3 mL). The organic layer was dried over anhydrous MgSO_4_, filtered and concentrated *in vacuo*. The residue was redissolved in CH_2_Cl_2_ (3 mL) and precipitated with a dropwise addition of Et_2_O to afford **Cmp1** as a beige powder (4.6 mg, 51%). The compound is unstable in solution under non-anhydrous conditions and decomposes to starting materials. The 10 mM stock of **Cmp1** in DMSO was immediately aliquoted and kept frozen at -20 °C. ^1^H NMR (400 MHz, DMSO-d_6_): δ 7.96 (1H, d, *J* = 2.7 Hz), 7.82 (1H, broad s), 7.61 (1H, d, *J* = 8.7 Hz), 7.47 (1H, dd, *J* = 8.4, 2.6 Hz), 7.32-7.14 (5H, m), 4.16-4.02 (1H, m), 3.07-2.81 (2H, m), 2.80-2.61 (2H, m), 2.38 (1H, d, *J* = 2.40 Hz), 1.28-1.16 (3H, m), 0.85-0.74 (1H, m).). LRMS (ES^+^): 475 ([^35^Cl^35^ClM+H]^+^, 100%), 477 ([^35^Cl^37^ClM+H]^+^, 75%), 479 ([^37^Cl^37^ClM+H]^+^, 20%). The compound has been characterized in the literature, data in agreement (Hoveyda et al., 2018).

### Plasmid constructions

FLAG-FFA2 was generated by amplification of mouse FFA2 plasmid using primers containing restriction sequences recognized by *Xbal* and *AfeI* and ligated with FLAG-LHR/pcDNA3.1 digested with *Xbal* and *AfeI* restriction sites. SEP-FFA2 was generated by subcloning SEP from SEP-LHR using *Xbal* and *AfeI* site and ligated to FFA2. All constructs used in the present study were verified by nucleotide sequence analysis.

### Cell Culture, transfections and stable cell lines

STC-1 and WT or β-ARR 1/2 KO HEK 293 cells were maintained in DMEM containing 10% FBS and penicillin/streptomycin (100 U/mL) at 37°C in 5% CO_2_. For both STC-1 or HEK 293 cells, transient transfections of plasmids were performed using Lipofectamine 2000 (Invitrogen) and cells were assayed 72 h post-transfection. Transfection of siRNA was performed using RNAiMAX (Invitrogen) and cells were assayed 96 h post-transfection. To generate FLAG-FFA2 and SEP-FFA2 stable cell lines, FLAG-or SEP-FFA2 was transfected in HEK 293 cells cultured in the presence of 0.5 µg/mL of geneticin.

### Signaling Assays

Intracellular cAMP and IP_1_ was determined by HTRF (cAMP Dynamic 2 and IP-one, respectively (CisBio)). For the measurement of intracellular cAMP, cells or colonic crypts were pre-treated with IBMX (0.5 mM, 5 min) prior to ligand stimulation (5 min). Measurement of IP_1_ was carried by incubating cells or colonic crypts with ligands in serum free media supplemented with 50 mM LiCL (2h). All cAMP and IP_1_ concentrations were corrected for protein levels. Calcium mobilization was measured by Fluo4-AM Direct Calcium Assay Kit (Invitrogen). Cells or colonic crypts were incubated with calcium dye in phenol red and serum free media for 30 min at 37°C and then at room temperature for 30 min. Cells or colonic crypts were imaged live using Leica SP5 confocal microscope using a 20X dry objective and a 488nm excitation laser. Movies were recorded at 1 frame per second for 1 min prior to ligand addition and a further 10-20 min following ligand addition to allow for calcium levels to lower to basal. For the measurement of p38 activation by western blot, STC-1 cells were serum-starved for 2 h prior to ligand stimulation. Following ligand stimulation, cells were rapidly washed in cold PBS and harvested with lysis buffer (1% Triton X-100, 50 mM Tris-HCl (pH 7.4), 150 mM NaCl, 0.5 mM EDTA, 1 mM NaF, 1 mM NaVO_3_ and a protease inhibitor tablet (Roche)). Cell extracts were separated on a 12% Tris-glycine polyacrylamide gel and transferred to a nitrocellulose membrane blotted with phospho-p38 MAPK antibody or p38 MAPK (Cell Signaling) as a loading control. Signal densities were quantified with ImageJ. Pre-treatments with either Ptx, YM-254890, Dyngo-4A or SB 203580 were carried out by incubating cells or colonic crypts for 20 h with 200 ng/mL Ptx, 5 min with 10 µM YM-254890, 45 min with 50 µM Dyngo-4a or 10 min with 5 μM SB 203580 before the addition of ligands. Experiments were conducted in duplicates for calcium mobilization and triplicates for all other experiments and were repeated at least three times.

### GLP-1 secretion assays

Plated STC-1 cells and colonic crypts were washed with secretion buffer (HBSS supplemented 1% BSA fatty acid free, which was adjusted to pH 7.4 with NaOH) and incubated in secretion buffer containing ligands for 2 h for STC-1 cells and 1 h for colonic crypts at 37°C. Inhibitors were used as for signaling assays. Following incubation, cell supernatants were collected, and the cells were lysed with lysis buffer (0.25 g sodium deoxycholate monohydrate, 0.88g NaCl, 0.5mL Igepal, 80 mM Tris HCL, pH 8, 1 tablet of complete EDTA-free protease cocktail inhibitor (Roche)). Samples were analyzed for GLP-1 secretion via an established in-house radioimmunoassay (Kreymann et al., 1987). The GLP-1 antibody has 100% cross-reactivity with all amidated forms of GLP-1 but does not cross-react with glycine extended forms. The intra-assay coefficients of variation for GLP-1 were 5.6%. As a control of GLP-1 release in the presence of inhibitors, cells or colonic crypts in the absence of inhibitors that secreted GLP-1 equivalent or less than NaCl were excluded from further analysis. All experiments were conducted in duplicate for colonic crypts and triplicate for STC-1 cells and repeated at least 3 times.

### Flow cytometry

Flow cytometry was used to quantitate internalization of FFA2 by measuring levels of receptor loss from the surface. Cells were fed live with M1 anti-FLAG antibody (20 min, 37°C) prior to treatment with ligands. Cells were then washed, lifted with PBS containing 2% FBS, centrifuged, and cell pellet washed with PBS and incubated with Alexa Fluor 488 secondary antibody (1h, 4°C). The fluorescence intensity of 10,000 cells were collected for each treatment and performed in triplicate using a FACS Calibur flow cytometer (BD Biosciences). Cells that were not exposed to any antibodies or secondary antibody alone were used for controls. All experiments were conducted at least three times.

### Immunofluorescence and confocal imaging

Receptor imaging in live or fixed cells were conducted by incubating live cells with FLAG M1-antibody (20 min, 37°C) and then with fluorescent secondary antibody (20 min, 37°C for live cell imaging) in phenol-red-free DMEM prior to agonist treatment. If inhibitors were used these were administered to the cells at appropriate time before ligand stimulation. To fix cells, cells were washed three times in PBS/0.04% EDTA to remove FLAG antibody bound to surface receptors prior to fixation with 4% paraformaldehyde in PBS (20 min), blocked with 2% FBS (1 h), permeabilized using 0.2% TritonX100, incubated with primary antibody (EEA1 or APPL1 (Cell Signaling) 1 h), washed and subsequently incubated with goat anti-mouse or rabbit Alexa Fluor secondary antibodies (Invitrogen) (1 h) at RT. Cells were washed again and mounted with Fluoromount-G (Thermo Fisher). Both live and fixed cells were visualized via a TCS-SP5 microscope (Leica) with a 63x oil-immersion objective and 1.4 numerical aperture (NA). Images were acquired using Leica LAS AF image acquisition software. Raw-image file were analyzed using ImageJ or LAS AF Lite (Leica) to measure endosomes diameter size or level of co-localization.

### TIRFM

Cells were imaged using the Elyra PS.1 AxioObsever Z1 motorized inverted microscope with a sCMOS or EMCCD camera and an alpha Plan-Aprochromat 100x/1.46 Oil DIC M27 Elyra objective (Zeiss), with solid-state lasers of 488 nm, 561 nm and/642 nm as light sources. For live cell imaging, approximately 15 minutes prior to imaging, culture media was replaced with Opti-MEM reduced serum media supplemented with HEPES. Imaging was then carried out using a Zeiss Elyra PS.1 microscope controlled at 37°C with 5% CO2. Time-lapse movies of whole cells were taken for 60 seconds, at 10 frames per second (fps) using Zen lite acquisition software. Fixed cells were prepared as for confocal imaging.

### Statistical analysis

Data are given as mean ± SEM. Mann-Whitney t-test, one-way ANOVA followed by Dunnett’s post-test, or two-way ANOVA followed by Bonferroni post-test was used when comparing two groups, more than two groups or at least two groups under multiple conditions, respectively. Statistical significance was determined using GraphPad Prism. The number of samples (n) has been indicated for each figure panel. Differences were considered significant p ≤ 0.05.

## ABBREVIATIONS

APPL1: adaptor protein containing
PH domain: PTB domain and leucine zipper motif;
B2AR: β2-adrenergic receptor;
Cmp1: compound 1;
DMEM: Dulbecco’s modified eagles medium;
EE: early endosome;
EEA1: early endosome antigen 1;
FFA2: free fatty acid receptor 2;
FFA3: free fatty acid receptor 3;
GAPDH: glyceraldehyde 3-phosphate dehydrogenase;
GPCR: G protein-coupled receptor;
LHR: luteinizing hormone receptor;
Ptx: pertussis toxin;
RIA: radio-immunoassay;
SEP: super ecliptic pHluorin;
SCFA: short chain fatty acid;
TIRFM: total internal reflection fluorescent microscopy;
VEE: very early endosome.

