## Supplementary figures and images for "Endosomal free fatty acid receptor 2 signaling is essential for propionate-induced anorectic gut hormone release"

### Supplemental figures

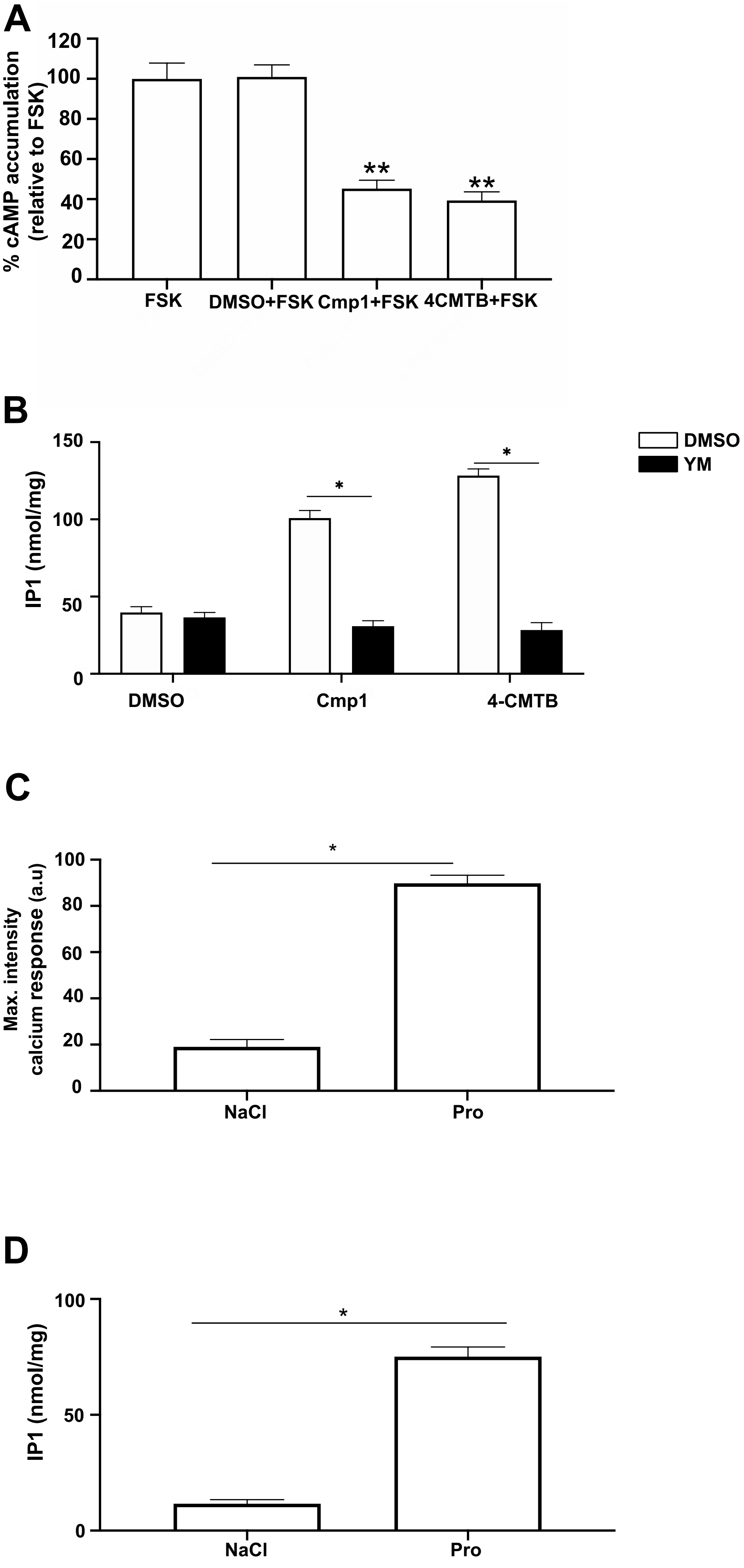

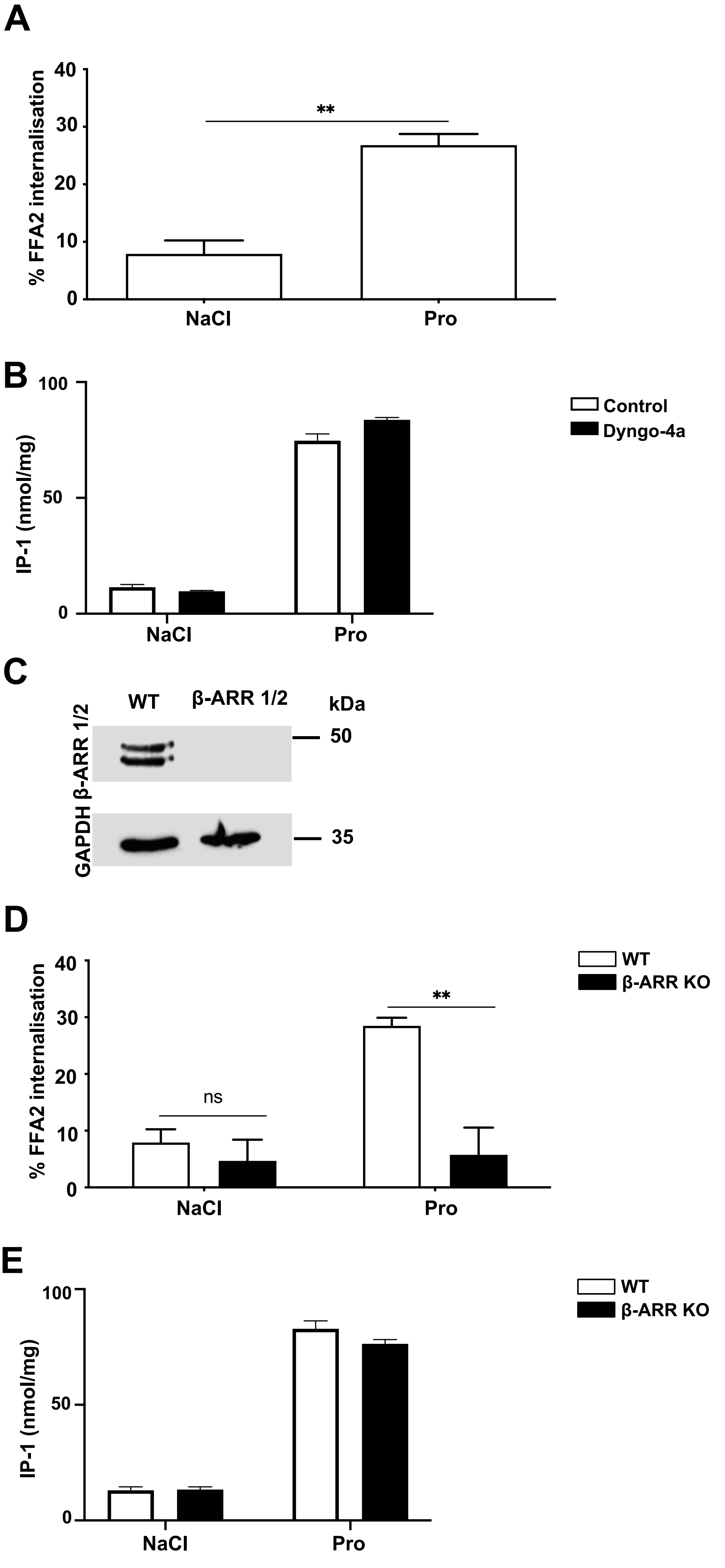

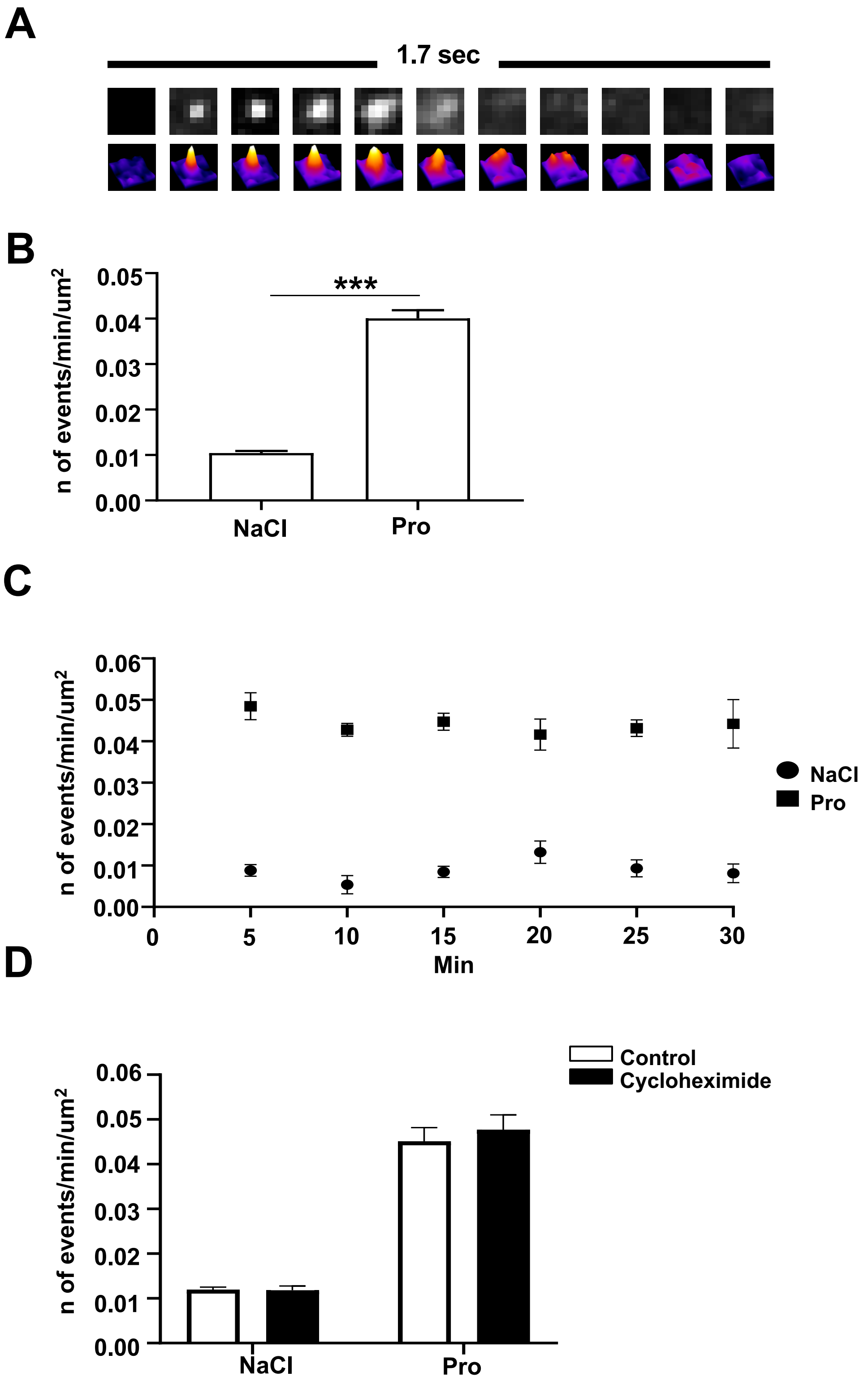

#### Figure S4

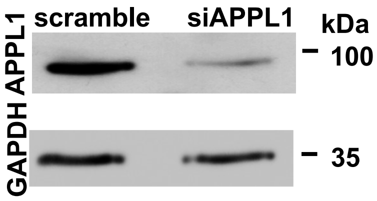

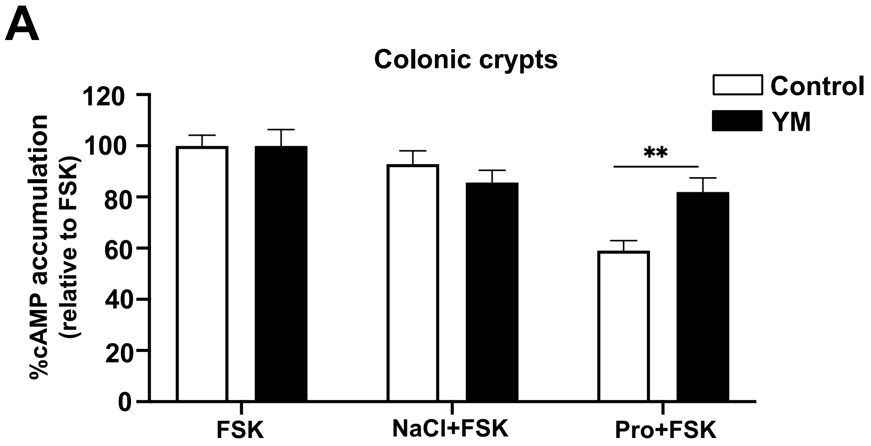

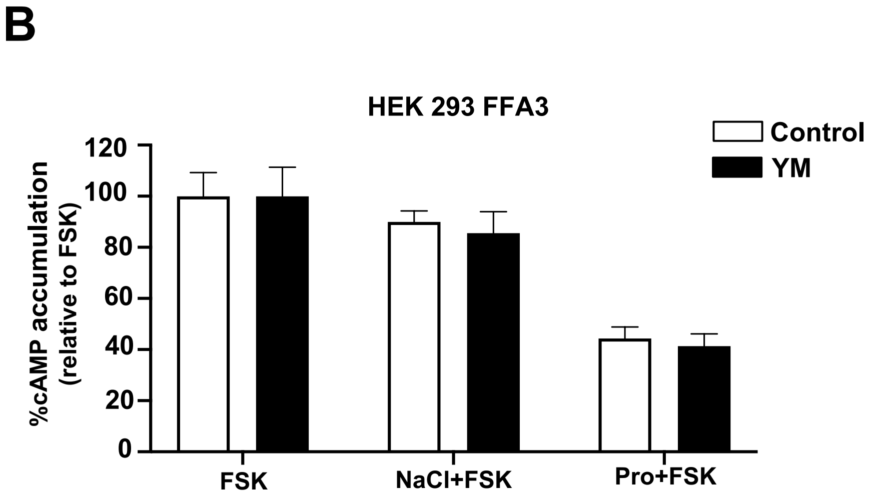

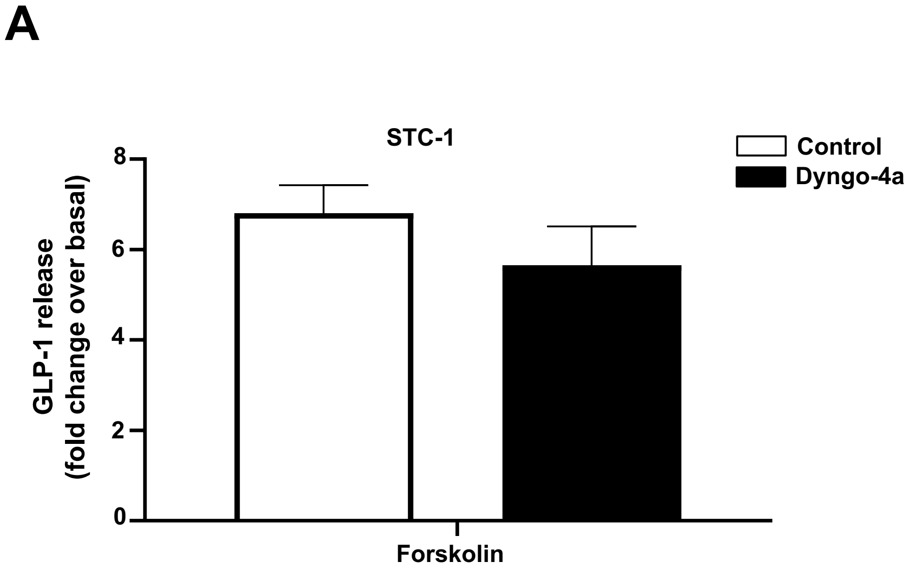

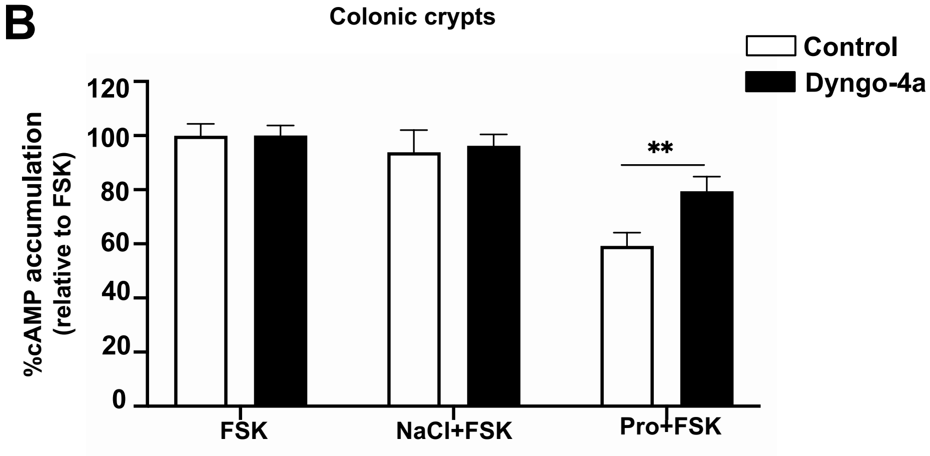
