## Supplemental figure legends for "Endosomal free fatty acid receptor 2 signaling is essential for propionate-induced anorectic gut hormone release"

**Figure S1 SCFAs are unable to signal via Gαq/11 in enteroendocrine cells.** (A) Gαi/o signaling activated by FFA2 ligands was measured via inhibition of forskolin (FSK)-induced cAMP signaling in STC-1 cells. Cells were stimulated with either DMSO (untreated control), Compound 1 (Cmp1) or 4-CMTB (10 µM, 5 min). n = 3 independent experiments. Two-sided Mann-Whitney U test, *** p < 0.001. (B) Intracellular IP_1_ levels measured in STC-1 cells, pre-treated with YM-254890 (YM, 10 µM, 5 min) and stimulated with either DMSO (untreated control), Compound 1 (Cmp1) or 4-CMTB (10 µM, 5 min). n = 4 independent experiments. Two-sided Mann-Whitney U test, *** p < 0.001. Intracellular calcium mobilization (C) or intracellular accumulation of IP_1_ (D) measured in HEK 293 cells expressing FLAG-FFA2. Cells were stimulated with either NaCl or sodium propionate (Pro) (1 mM). For intracellular calcium mobilization, average maximal intensities of n = 20 cells in duplicate per 4 independent experiments. Two-sided Mann-Whitney U test, * p < 0.05. For IP_1_, n = 4 independent experiments. Two-sided Mann-Whitney U test, *** p < 0.00. Data represent mean ± SEM.

**Figure S2 Differential spatial requirement for FFA2-mediated signaling, related to Figure 2.** (A) FFA2 internalization measured by flow cytometry. HEK 293 cells transiently expressing FLAG-FFA2 were treated with and without NaCl (1 mM), or sodium propionate (Pro) (1 mM, 20 mins). FFA2 internalization was determined by decrease cell-surface labeling compared to untreated control. n = 4 independent experiments. Two-sided Mann-Whitney U test, ** p < 0.01. (B) Intracellular accumulation of IP_1_ measured in HEK 293 cells transiently expressing FLAG-FFA2 pre-treated with either DMSO (vehicle) or Dyngo-4a (50 µM, 45 min) and treated with either NaCl or propionate (Pro) (1 mM, 2 h). n = 24 independently plated wells for either control or in the presence of Dyngo-4a, representative of 4 independent experiments. Two-sided Mann-Whitney U test. (C) Western blot confirmation of the absence of β-arrestin 1/2 (β-ARR) expression in β-ARR KO HEK 293 cells compared to WT cells. GAPDH was used as loading control. (D) Dependence of FFA2 internalization on β-ARR measured by flow cytometry. WT and β-ARR KO HEK 293 cells transiently expressing FLAG-FFA2 were treated and data analysis were carried as in (A). n = 4 independent experiments. Two-sided Mann-Whitney U test, ** p < 0.01. (E) Intracellular accumulation of IP_1_ measured in WT or β-ARR KO transiently expressing FLAG-FFA2 cells and treated as in (B). n = 4 independent experiments for either WT or β-ARR KO transiently expressing FLAG-FFA2. Two-sided Mann-Whitney U test. Data represent mean ± SEM.

**Figure S3 Characterization of SEP-FFA2 recycling events via TIRFM, related to Figure 4C.** (A) Series of TIRFM images of a single SEP-FFA2 recycling event following propionate stimulation with surface plot of fluorescence. (B) Number of recycling events measured by TIRFM in HEK 293 cells stably expressing SEP-FFA2 following stimulation of NaCl (1 mM) or propionate (Pro, 1 mM); n = 15 cells per condition; collected across 3 independent experiments. Two-sided Mann-Whitney U test, *** p < 0.001. (C) Number of recycling events over time following stimulation of NaCl (1 mM) or propionate (Pro, 1 mM) measured by TIRFM in HEK 293 cells stably expressing SEP-FFA2; n = 4 cells per time point; collected across 3 independent experiments. (D) Number of recycling events measured by TIRFM in HEK 293 cells stably expressing SEP-FFA2, pre-treated with DMSO (vehicle) or cycloheximide (10 μg/mL, 90 minutes) prior to NaCl (1 mM) or propionate (1 mM) stimulation. n = 10 cells per condition, collected across 3 independent experiments. Two-sided Mann-Whitney U test. Data represent mean ± SEM.

**Figure S4 APPL1 knockdown in STC-1 cells via siRNA, related to Figure 4D.** Representative western blot of total cellular levels of APPL1 in lysates collected from STC-1 following either scramble (control), APPL1 (siAPPL1) siRNA-mediated knockdown. GAPDH was used as loading control.

**Figure S5 The Gαq/11 inhibitor, YM-254890, partially inhibits Gαi/o signaling activated by propionate/FFA2.** (A) Gαi/o signaling activated by propionate (Pro, 1 mM, 5 min) in colonic crypts was measured via inhibition of forskolin (FSK)-induced cAMP signaling and pre-treated with or without YM-254890 (YM, 5 min, 10 μM). n = 4 independent experiments. Two-sided Mann-Whitney U test, **, p < 0.01. (B) HEK 293 cells transiently expressing FFA3, was pre-treated with and without YM, and Gαi signaling measured as in (A). n = 4 independent experiments. Two-sided Mann-Whitney U test, **, p < 0.01. Data represent mean ± SEM.

**Figure S6 Dyngo-4 partially inhibits Gαi/o signaling activated by propionate/FFA2 in colonic crypts** (A) Forskolin induced GLP-1 release from STC-1 cells in the presence of Dygno-4a. STC-1 cells were pre-treated with either DMSO or Dyngo-4a (50 µM, 45 min). n = 5 independent experiments. Two-sided Mann-Whitney U test. (B) Gαi/o signaling activated by propionate (Pro, 1 mM, 5 min) in colonic crypts was measured via inhibition of forskolin (FSK)-induced cAMP signaling and pre-treated with or without Dygno-4a (50 µM, 45 min). n = 4 independent experiments. Two-sided Mann-Whitney U test, **, p < 0.01. Data represent mean ± SEM.
